# Development of a new cold hardiness prediction model for grapevine using phased integration of acclimation and deacclimation responses

**DOI:** 10.1101/2022.09.09.507298

**Authors:** Al P. Kovaleski, Michael G. North, Timothy E. Martinson, Jason P. Londo

## Abstract

Cold injury limits distribution of perennial agricultural crops, though replacement of plants and other management practices may allow for some damage tolerance. However, winter damage to crops such as grapevines (*Vitis* spp.) can result in losses in yield the following year if buds are damaged, but over many years when vines must be replaced and reach maturity before fruiting. Despite risks, grapevines are cultivated at the edge of permissible climate and rely on cold hardiness monitoring programs to determine when cold damage mitigation and management practices are required. These monitoring programs represent a critical, but laborious process for tracking cold hardiness. To reduce the need for continuous monitoring, a model (WAUS.2) using cold hardiness data collected over many years from Washington state, USA, growers was published in 2014. Although the WAUS.2 model works well regionally, it underperforms in other regions. Therefore, the objective of this work was to develop a new model (NYUS.1) that incorporates recent knowledge of cold hardiness dynamics for better prediction outcomes. Cold hardiness data from *V. labruscana* ‘Concord’, and *V. vinifera* ‘Cabernet Sauvignon’ and ‘Riesling’ from Geneva, NY, USA were used. Data were separated in calibration (~2/3) and validation (~1/3) datasets. The proposed model uses three functions to describe acclimation, and two functions to describe deacclimation, with a total of nine optimized parameters. A shared chill response between acclimation and deacclimation provides a phased integration where acclimation responses decrease over the course of winter and are overcome by deacclimation. The NYUS.1 model outperforms the WAUS.2 model, reducing RMSE by up to 37% depending on cultivar. The NYUS.1 model also tends to be more conservative in its prediction, slightly underpredicting cold hardiness, as opposed to the overprediction from the WAUS.2 model. Some optimized parameters were shared between cultivars, suggesting conserved physiology was captured by the new model.

**Highlights:**

- Multi-year cold hardiness data from three grapevine cultivars were used for modeling
- Cold hardiness was modeled based on daily temperature and accumulated chill
- Phased acclimation and deacclimation processes result in cold hardiness predictions
- The new model was compared to the currently available model for grapevines
- The model proposed here outperforms the currently available model

## 1. Introduction

One of the most important abiotic stresses limiting the distribution of plants worldwide is low winter temperatures. Single freezing events can effectively eliminate plant populations, thus limiting the cold range edge of native plants (Körner, 2021). Perennial agricultural plants are often introduced beyond their native cold range limits and therefore have higher risk of cold injury during winter (Hänninen, 2013). This is the case for grapevines (*Vitis* spp.), especially domesticated European grapevines (*V. vinifera*), in northern regions within the United States and in Canada, where low temperatures during the dormant season in winter are a limiting factor. In these agricultural systems, injured or dead plants can be replaced, although this increases production costs and can create gaps or delays in production. Therefore, understanding and predicting cold hardiness of perennial plants can be helpful in determining management practices and site selection based on winter survival.

Domesticated European grapevines are native to the Mediterranean yet are cultivated on all continents except Antarctica. Within the US, New York state is the third largest grape producer in the US (USDA, 2019), growing grapevines in significantly colder conditions than that of the larger two producing states (California and Washington), and colder than some regions with which it shares latitudes within Europe, such as northern Spain, southern France, and central Italy. Viticulture in New York relies on the proximity to large bodies of water (e.g., Lake Erie, Lake Ontario, the Finger Lakes, Atlantic Ocean) to help buffer low temperatures during winter, thus allowing cultivation of *V. vinifera* grapevines, which are generally less cold hardy than American species (e.g., *V. labrusca*, *V. riparia*) and hybrid varieties (Londo and Kovaleski, 2017). Winter survival of mixed buds in grapevines is required for production of grapes in the following growing season. These buds are formed during summer and must remain dormant and develop cold hardiness throughout fall to survive low temperatures in winter before resuming growth in spring. The ability to predict bud cold hardiness would allow growers to monitor conditions and decide when it may be necessary to evaluate winter bud injury, particularly in currently cold growing regions, but accurate predictions could also be used to evaluate current suitability for expansion into other areas, or for identifying regional suitability changes as climate changes. While climate predictions suggest winter temperatures will rise, more erratic climate is also expected with large swings between warm temperatures and acute low temperatures (IPCC, 2022). As cold damage mitigation tools are developed, accurate predictions of cold hardiness are also needed to assess necessity and timing of deployment of these methods (Li and Dami, 2016; Kovaleski and Londo, 2019). Therefore, accurate cold hardiness models can help evaluate the future of viticulture in cold climates from a winter survival standpoint.

Dormancy and cold hardiness are separate but concomitant processes within a plant (Arora et al., 2003; Kovaleski, 2022). Both processes are required for survival of low temperatures during the cold season in subtropical and temperate environments but appear to respond to slightly different environmental cues. Dormancy establishment in grapevines typically occurs in response to shortening photoperiod in late summer and fall (Fennel and Hoover, 1991) but other environmental factors can also play a role [e.g., drought (Shellie et al., 2018)]. Dormancy progression is then modulated by exposure to low temperatures (chilling), transitioning from a warm temperature non-responsive (endodormancy) to a responsive phase (ecodormancy), from which plants are released from upon return of warm temperatures in spring (Lang et al., 1987).

Cold hardiness usually presents a U-shape pattern during the progression from fall to winter to spring with three distinct phases (Ferguson et al., 2011; Ferguson et al., 2014; Cragin et al., 2017; Londo and Kovaleski, 2017; Kovaleski, 2022). In the fall, acclimation leads to gains in cold hardiness of grapevine buds through the establishment and enhancement of supercooling. In mid-winter, buds reach maximum cold hardiness and maintain high levels of cold hardiness depending on environmental cues. Finally, in late winter and early spring as warm temperatures return, deacclimation leads to loss of cold hardiness – though reacclimation is possible upon return of low temperatures. Photoperiod alone is unable to elicit significant gains in cold hardiness (Fennel and Hoover, 1991). While some authors have indicated that only the synergistic effect of short photoperiod and low temperatures can induce deep cold hardiness (Schnabel and Wample 1987), temperature effects appear to be the largest contributors to cold hardiness levels of grapevine buds, and models solely based on temperature have shown to produce good regional predictions (Ferguson et al., 2011; Ferguson et al., 2014; North et al., 2021).

Dormancy directly affects cold hardiness dynamics. During the endodormant period, in the fall and early winter, buds respond to low temperatures to gain cold hardiness. The ability to gain cold hardiness appears to decrease as chilling accumulates and buds transition to ecodormancy (Ferguson et al., 2011; Ferguson et al., 2014; North et al., 2021). The inverse seems to happen with regards to warm temperatures: a bud’s deacclimation responsiveness increases as chilling accumulates (Ferguson et al. 2011, 2014; Cragin et al., 2017; Kovaleski et al., 2018; North et al., 2021; Kovaleski, 2022). Therefore, acknowledging dormancy dynamics is necessary for predicting cold hardiness.

Currently, the most widely used model for predicting grapevine bud cold hardiness was developed using cold hardiness data from Washington state based on *V. vinifera* ‘Cabernet Sauvignon’, ‘Chardonnay’, and *V. labruscana* ‘Concord’ (Ferguson et al., 2011), and was later enhanced and further expanded to add twenty other cultivars (Ferguson et al., 2014). Here we propose a new nomenclature for these models, using the region and country of development, as well as a version number. Therefore, from here on the models proposed by Ferguson and collaborators are referred to as WAUS.1 (Ferguson et al., 2011) and WAUS.2 (Ferguson et al., 2014). The WAUS.2 model has also been recently adapted for cold climate interspecific hybrids grown in a different environment (North et al., 2021). However, as indicated by the original authors, local adaptation of the model may be required given the specific climate in which it was developed. Indeed, the model does not produce reliable results in viticultural regions in the Eastern United states, including the state of New York. This is likely a consequence of trade-offs among three dimensions in models: accuracy, realism, and generality (Levins, 1966; Levins, 1968). Accuracy represents the fit between predictions and observations, realism refers to causally appropriate internal model processes, and generality represents robust applicability across space and/or time (Kramer et al., 2002). A model will ideally maximize all accuracy, realism, and generality simultaneously; however, in practice, model developers must identify an optimal point among the three dimensions that matches the overall aim of a model (Hänninen, 2016). In the case of the WAUS.2 model, accuracy was prioritized over realism and generality. Recent studies of dormancy related to cold hardiness have shown that some of the parameters used in the model, such as the linear response to temperature and the discrete change in responses by the vines based on dormancy status do not accurately describe the nature of the process (Kovaleski et al., 2018; North et al., 2022). In fact, linear responses to temperature have been related to errors in predictions of spring phenology as well (Wolkovich et al., 2021).

The objective of this study was to develop a new model (NYUS.1) for cold hardiness prediction of grapevine buds with greater accuracy for the Eastern United States and other viticultural regions more broadly. To do so, we use a medium-term dataset comprised of three cultivars: *V. labruscana* ‘Concord’, and *V. vinifera* ‘Cabernet Sauvignon’ and ‘Riesling’. The choice of cultivars was made based on availability of data (partially due to regional relevance), and variability in responses. The new model expands from previously empirically determined concepts of deacclimation in response to chilling and forcing temperatures (Kovaleski et al., 2018; Kovaleski, 2022; North et al., 2022), thus with increased realism, and incorporates theoretical derivation of acclimation functions based on preliminary experimental data to predict cold hardiness with field-based temperature data.

## 2. Materials and Methods

### 2.1. Cold hardiness data

Bud cold hardiness data from three cultivars were used: *V. labruscana* ‘Concord’, *V. vinifera* ‘Cabernet Sauvignon’, and *V. vinifera* ‘Riesling’ (from here on referred to by cultivar name only). Buds were collected from vineyards located in Geneva, NY, throughout multiple dormant seasons in varying numbers depending on cultivar (for details, see **Supplementary Table S1**). Cold hardiness was measured using differential thermal analysis (DTA) (Wolf and Pool, 1987; Mills et al., 2006; Ferguson et al., 2011; Ferguson et al., 2014; Londo and Kovaleski, 2017, North et al., 2021). Briefly, for DTA, buds are placed on thermoelectric modules connected to a datalogger and placed into a programmable freezer which can gradually lower temperature over time. Once intracellular water freezes, heat release is recorded as a low temperature exotherm (LTE) and signifies the temperature of death for any given bud (see Mills et al., 2006 for additional details). The number of buds used to measure cold hardiness for each cultivar on each sampling date varied between 5 and 30 buds (although number of recovered peaks may have been lower). For model development, average LTE values for each sampling date were used.

Eight seasons of data are available for ‘Concord’, five for ‘Cabernet Sauvignon’, and nine for ‘Riesling’. The number of observations in each season varied based on cultivar, length of the season, and research program. For some seasons, more than one program collected data independently, but data were combined for modeling without using lab as a factor.

### 2.2. Weather data

Weather data were obtained from the Network for Environment and Weather Applications (NEWA; www.newa.cornell.edu). Hourly data were downloaded from which daily maximums and minimums were extracted for model development.

### 2.3. Data analyses

All data were analyzed with R (ver. 4.1.2; R Core Team, 2021) within R Studio (RStudio Team, 2021). Chill accumulation was calculated from hourly temperature data for each winter based on the dynamic model (chill portions; Fishman et al., 1987a; Fishman et al., 1987b) using the chillR package (Luedeling, 2020). Three inputs were then used for model development, daily maximum (*T_max_*) and minimum temperatures (*T_min_*), and chill accumulation (*chill*, which is a function of temperature).

### 2.4. Model development

The model presented here is based on a generalized function for cold hardiness (*CH*) from Kovaleski (2022) where the *CH* at any time *t_i_* during the dormant season is a result of the integration of separate acclimation and deacclimation processes occurring starting from bud set to *t_i_*:

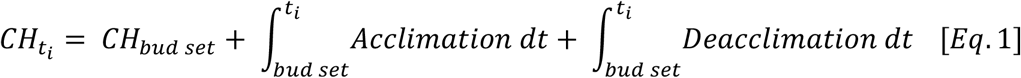

where *CH_bud set_* is the cold hardiness of buds as they set and mature in late summer and fall, which is a characteristic of the tissues, and *Acclimation* and *Deacclimation* are comprised of functions described below. Here, *Acclimation* and *Deacclimation* are integrated over daily time steps for *t*, but both are functions of temperature, either directly or indirectly through *chill*. However, field temperatures are themselves a function of time.

#### 2.4.1. For *Acclimation*

three functions were used and are based on expectations of the responses (visual representations of these are presented in **Figs. 1A-D**). First, a function is used to describe what is the maximum temperature below which any acclimation occurs as a response to chilling accumulation [the acclimation temperature threshold 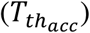]:

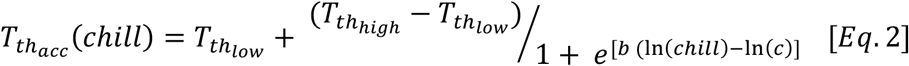

where the temperature threshold for acclimation decreases from a high threshold 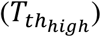 at low chill accumulations to a low threshold 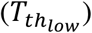 at high chill accumulations; *b* is the slope associated with how quickly the decrease occurs, and *c* is the inflection point of the log logistic curve (**Fig. 1A**). This decrease in temperature threshold represents physiological changes during chill accumulation and dormancy progression affecting the directionality of the cold hardiness dynamics process: at high chill accumulations, the temperature experienced by buds must be lower to promote any acclimation. Both parameters *b* and *c* are shared with deacclimation potential, which is explained further below (in ***2.4.2. Deacclimation***).

**Figure 1.**
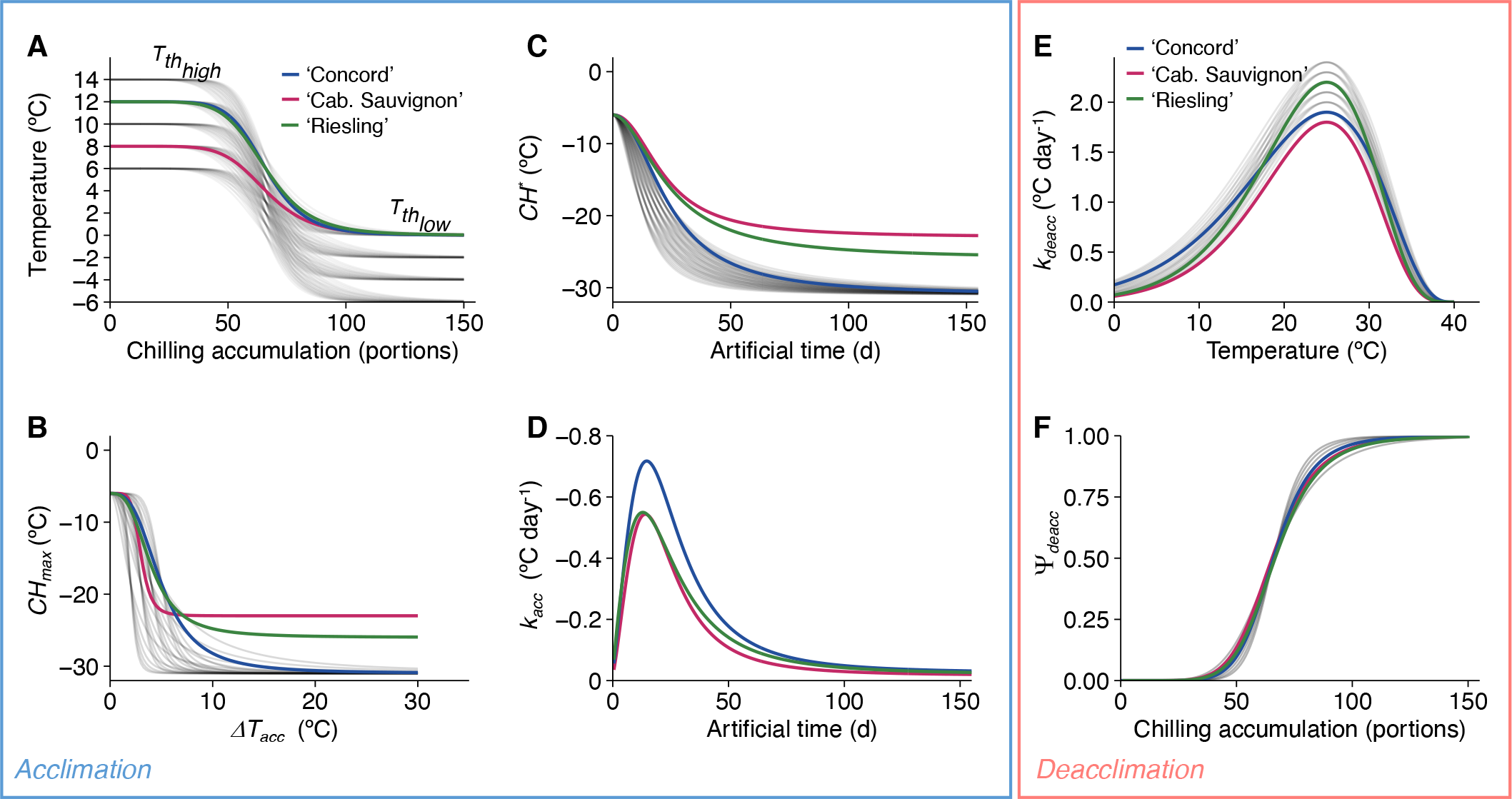
Graphic representation of functions used to model cold hardiness of grapevine buds for ‘Concord’, ‘Cabernet Sauvignon’, and ‘Riesling’. Functions are separated between acclimation (**A-D**) and deacclimation (**E, F**). Solid colored lines represent fits (**A, B, E, F**), or example fits for each cultivar (**C, D**); blue = ‘Concord’, magenta = ‘Cabernet Sauvignon’, green = “Riesling’. The semi-transparent grey lines represent iterations of combinations for ‘Concord’ parameters that were not selected. **(A)** Acclimation step 1; see also **Eq. 2**. The temperature below which acclimation occurs is the acclimation temperature threshold 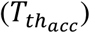, which changes according to chill accumulation. As chill accumulates, 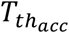 changes from high acclimation temperature threshold 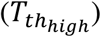 to low acclimation temperature threshold 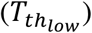. The driver of acclimation, *DT*, is calculated from the difference between the daily low temperature and 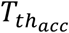. **(B)** Acclimation step 2; see also **Eq. 3**. Upper limit of curves, or maximum cold hardiness (*CH_max_*) at *DT_acc_* = 0°C, represents cold hardiness at bud set (*CH_bud set_*), which was fixed at –6 °C for all cultivars. Lower limit of curves, or *CH_ax_* at *DT_acc_* = 30°C, represents the maximum cold hardiness observed during any season (*CH_abs max_*), which we set equal to the average of the 15 lowest cold hardiness measurements rounded down to the nearest integer. The shape of these curves between the upper and lower limits are determined by fitting the slope (*d*) and intercept (*f*) parameters in **Eq. 3**. **(C)** Acclimation step 3; see also **Eq. 4**. Example curves for artificial cold hardiness (*CH^*^*) for a maximized *DT* in response to artificial time. These curves represent the time buds would take when exposed to a given *DT_acc_* to reach the *CH_max_* elicited. **(D)** Final acclimation rate based on partial derivative of step 3; see also **Eq. 5**. The acclimation rate *k_acc_* is obtained based on the artificial time calculated from Eq. 4 solved for time using the the cold hardiness of the previous day as *CH^*^*. **I** Deacclimation step 1, deacclimation rate; see also **Eq. 6**. The deacclimation rate (*k_deacc_*) varies across temperature from 0 °C to 40°C. For all cultivars, *k_deacc_* reaches maximum at 25 °C (*T_optim_*) and is zero at 40 °C (*T_max_*). Two aspects that were fit separately for each cultivar are the maximum deacclimation rate 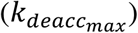 and the coefficient *a*, which determines the slope on either side of 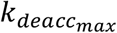. **(F)** Deacclimation step 2, deacclimation potential; see also **Eq. 7**. Deacclimation potential (*Ψ_deacc_*) modulates the deacclimation rate to an effective deacclimation rate 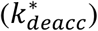 based on chill accumulation from low to high (i.e., *Ψ_deacc_* increases as chill accumulates). The inflection point for all cultivars was fixed at 66 chill portions, while the slope parameter of *Ψ_deacc_* curves (*b*) was fit separately for each cultivar and is shared with 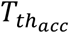.

The second function describes the maximum cold hardiness possible (*CH_max_*) elicited by the difference between the *T_min_* of any given day and the 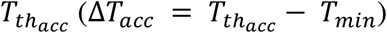. Here, we used a log-logistic function to describe the response:

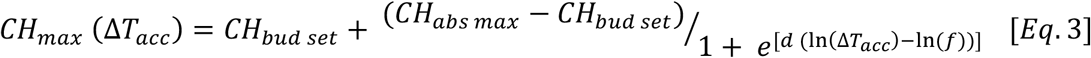

where *CH_bud set_* is the minimum cold hardiness, which was conservatively fixed at –6 °C for all cultivars (the resolution of the limit of detection of our equipment); *CH_abs max_* is the maximum cold hardiness observed during any season, and here was used as the average of the 15 lowest cold hardiness measurements rounded down to the nearest integer; *d* is the slope associated with the logistic curve; and *f* is the inflection point of the logistic curve (**Fig. 1B**).

The third function describes the rate of acclimation based on how close to the *CH_max_* the cold hardiness of the previous day 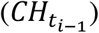 is. For simplicity, here we present this in two steps. First, the shift from *CH_bud set_* to the *CH_max_* elicited by any given *ΔT_acc_* is not instant. Should buds be exposed to a constant *ΔT_acc_*, acclimation would occur over time, giving rise to rate of acclimation. Because these are based on assumptions related to acclimation rather than experiments, we have dubbed this cold hardiness as an artificial cold hardiness (*CH^*^*). The artificial cold hardiness (*CH^*^*) thus changes from *CH_bud set_* to *CH_max_* as a function of time based on a log-logistic function:

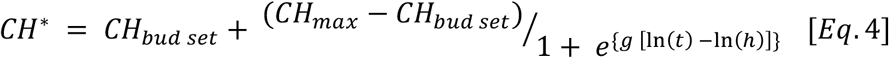

where *g* and *h* are the slope and intercept associated with a log-logistic curve (**Fig. 1C**). *CH^*^* is also indirectly a function of the temperature and chill accumulation since both contribute to defining *CH_max_*. An artificial time *t_a_* is found by solving **Eq. 4** for *t* using 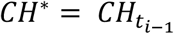 (the cold hardiness of the previous day) and *CH_max_* for *T_min_* at *t_i_* (using **Eq. 3**). Using *t_a_*, we can obtain the rate of acclimation (*k_acc_*) in the field at any time *t_i_* through the partial derivative of the log logistic function with respect to time:

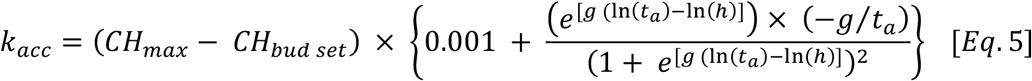

Where *g* and *h* are the slope and intercept associated with a log-logistic curve, and *t_a_* is the artificial time based on 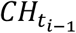 (**Fig. 1D**). Here, *k_acc_* is a function of artificial time (since field temperature changes continuously), and the field temperature and accumulated chill that define *CH_max_*. In addition to the rate obtained through the partial derivative, a 0.001 (or 0.1%) rate was added to acclimation at all times. This was incorporated to acknowledge anecdotal observations that buds seem to continue acclimating when temperatures are very low (Kovaleski personal observations). Therefore, buds can surpass the *CH_max_* elicited by any *ΔT_acc_*, and even a cultivar’s *CH_abs max_*, if temperatures are continuously low. While the actual effect of the factor is very small in this model, the factor could be enhanced or reduced to accommodate new perspectives based on future research. A sample workflow of an *Acclimation* step is shown in **Fig. 2**.

**Figure 2.**
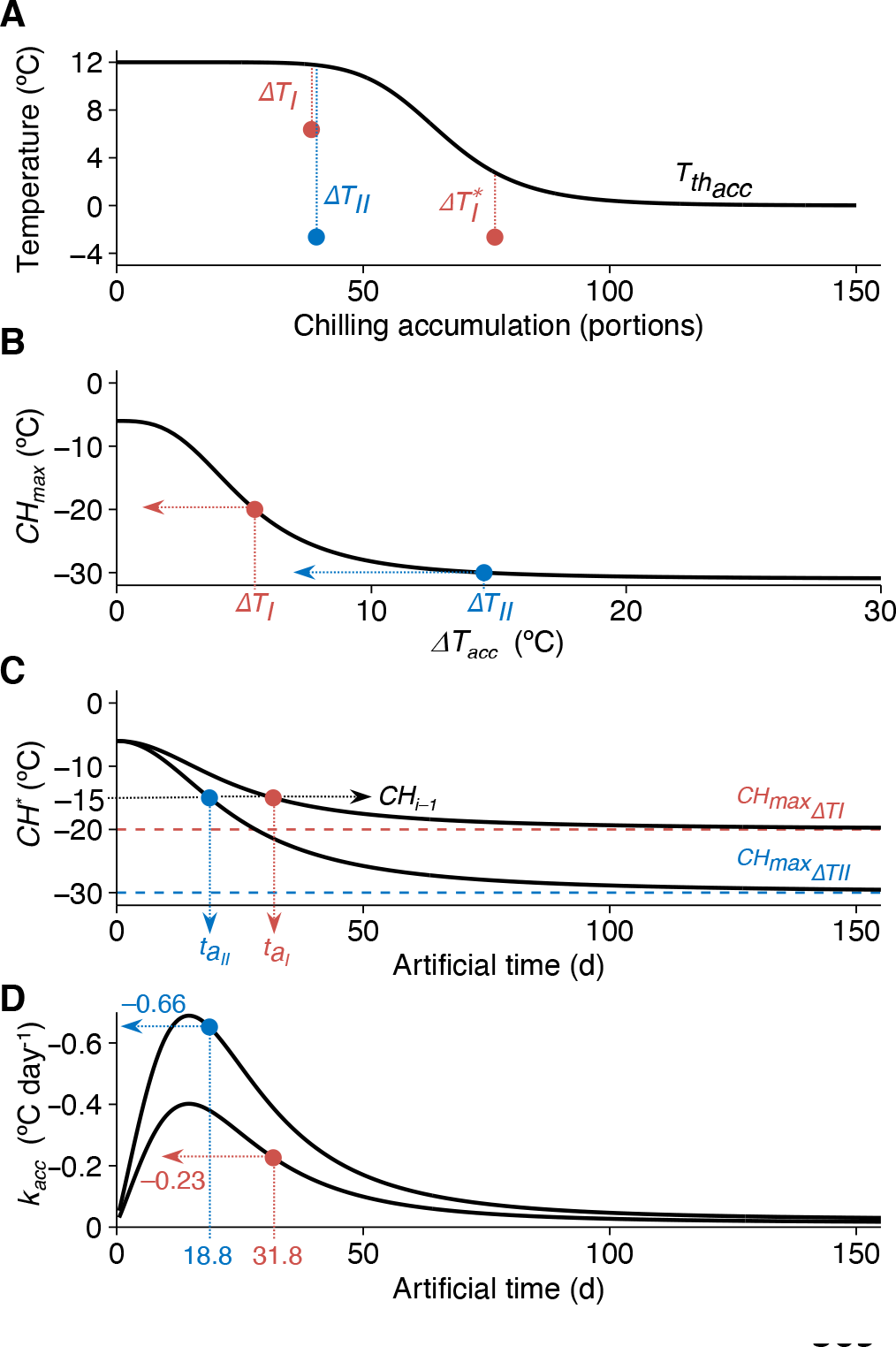
Example workflow to describe progression for acclimation step in the NYUS.1 model and highlight relative importance of changes in conditions. **(A)** The 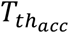 in this example varies from 12 °C 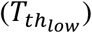 to 0 °C 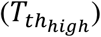 across accumulating chill. *DT* is the difference between *T_min_* (points) and 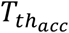. Two *DT_acc_* scenarios are included in this example (*DT_I_* and *DT_II_*) to highlight importance of conditional changes throughout acclimation calculations. Here we observe that different temperatures can lead to the same *DT* (*DT_I_* = *DT^*^_I_*): as chill accumulates and 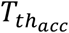 decreases, a lower *T_min_* would elicit the same response. We can also observe that the same temperature can lead to different *DTs* (*DT_II_* ≠ *DT^*^_I_*) if at two different chilling accumulations. **(B)** Based on a given *DT_acc_*, a certain *CH_max_* is obtained, which is greater (in magnitude) for greater *DT_acc_*. **(C)** With the different *CH_max_* for each of the scenarios, *DT_I_* and *DT_II_*, we can observe the different curves for gains in cold hardiness if buds were constantly exposed to those temperatures. Solving ***Eq. 4*** for time using the *CH* of the previous day (*CH_i-1_*), here –15 °C, as *CH^*^* we obtain the time at which this would occur along that artificial temperature exposure: 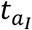 and 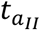. **(D)** Using the first derivative of those acclimation curves, we obtain the curves for rate of acclimation, *k_acc_*. The rate of acclimation for a given day, *t_i_*, is calculated by solving ***Eq. 5*** using 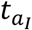 and 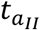.

#### 2.4.2. For *Deacclimation*

two functions were used (**Figs. 1E, F**) which are based on descriptions from previous empirical experiments on grapevines and other species (Kovaleski et al., 2018; Kovaleski, 2022; North et al., 2022). One is for the rate of deacclimation (*k_deacc_*) that any given temperature elicits. Other functions have been used to describe *k_deacc_*, such as a logistic (Chuine, 2000; North et al., 2022), combination of exponential and logarithmic (Kovaleski et al. 2018) or a third-degree polynomial (Kovaleski, 2022). Here, the temperature response curve used was that based on a temperature-photosynthesis curve (O’Neil et al., 1972):

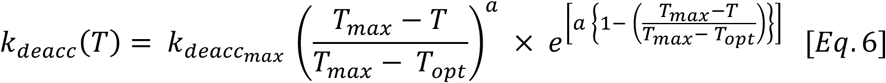

where *k_deacc_* (*T*) is the rate of deacclimation (°C day^−1^) at temperature *T*(°C); *a* is a coefficient related to the slope of the curve; 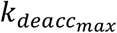 is the maximum rate of deacclimation at the optimal temperature (*T_optim_*, °C), and *T_max_* is the maximum temperature (°C) from which point the deacclimation rate becomes zero. *T_optim_* and *T_max_* were fixed at 25 °C and 40 °C, respectively. While an absolute decrease in rates with temperature has not observed in empirical studies (Kovaleski et al., 2018; North et al., 2022), some considerations can be made: a deviation from exponential increase in rates was observed (Kovaleski et al., 2018; Kovaleski, 2022); temperatures much higher than 25 °C do not occur frequently during the deacclimation period in the spring; and the suggestion of deacclimation as an enzymatic process is acknowledged by this curve (Kalberer, 2006; Kovaleski et al., 2018). The *T_optim_* of 25 °C was used based on previously reported deacclimation rates (Kovaleski et al., 2018; North et al., 2022). Because of the non-linearity of the response, the amount of deacclimation observed in a day was the average between *k_deacc_* for the daily *T_max_* and *T_min_*, rather than using the average of the temperature.

The second function defining deacclimation is the deacclimation potential (*Ψ_deacc_*), which modulates the level of effective deacclimation occurring based on a log logistic function of chill accumulation during the winter:

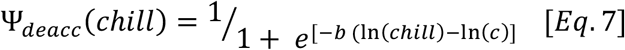

where *b* is the slope associated with the log-logistic curve, and *c* is the inflection point of the log-logistic curve, a chilling requirement-analogous measurement where 50% of the maximum *Ψ_deacc_* is achieved. Because *b* and *c* were shared with 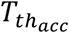, such that the dormancy-related responses to *chill* are symmetric for *Acclimation* and *Deacclimation*, a negative sign is used for *b* here (negative values indicate increases in *Ψ_deacc_* as chill accumulation increases). For *c*, a fixed value of 66 chill portions was used based on previous work with many species (Kovaleski, 2022). The phased integration of *Acclimation* and *Deacclimation* provided by the shared *b* and *c* terms mean that as chilling accumulates, *Acclimation* becomes more difficult, requiring lower temperatures, while *Deacclimation* becomes more pronounced.

The effective deacclimation rate at any time *t_i_*, 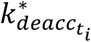, is ultimately a result of the product of the average daily deacclimation rate and the deacclimation potential:

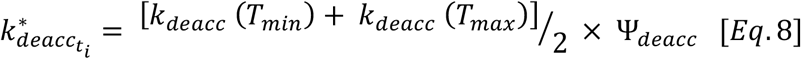

### 2.5. Model optimization

The cold hardiness data available was separated in calibration and validation datasets by season-year. The ratio was approximately 2/3 calibration and 1/3 validation. The years were randomly assigned to each dataset within each cultivar using set.seed() based on when code preparation for each cultivar started [17 August for ‘Concord’ set.seed(1708); 7 June for ‘Cabernet Sauvignon’ set.seed(0706); 7 July for ‘Riesling’ set.seed(0708)]. The dynamic model based on **Eq. 1** predicts cold hardiness with 1-day time steps. Nine parameters were optimized here: 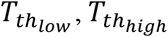, *b*, *d*, *f*, *g*, *h*, 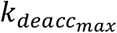, and *a* (see **Table 1** and **Fig. 1**). The parameters were optimized using a stepwise iterative method based on the minimization of root mean square error (RMSE) between predicted and observed CH for a calibration dataset (RMSE_cal_). The parameter combination with lowest RMSE_cal_ was then used to predict CH using a naïve validation dataset, and RMSE of a prediction was then calculated (RMSE_val_). The RMSE_val_ of the present model was compared to that of the predictions based on the latest update of the currently available grapevine bud cold hardiness developed in Washington state, WAUS.2 (RMSE_WA_; Ferguson et al., 2014 – see also Ferguson et al., 2011 for details on equations). Other fitness statistics were also calculated for both models [*Bias* = *CH_measured_* − *CH_predicted_*; Pearson correlation (*r*)].

**Table 1.** Parameter levels tested in all combinations stepwise for three grapevine cultivars for each of six equations. A total of 3,386,880 combinations were tested per cultivar for all years in the calibration datasets.

| Cultivar | $T_{thlow}$ | $T_{thhigh}$ | $b$ | $c$ | $CH_{abs\ max}$ | | | $CH_{bud\ set}$ | $d$ | $f$ | $g$ | $h$ | $k_{deacc}$ | | | $a$ |
| --- | --- | --- | --- | --- | --- | --- | --- | --- | --- | --- | --- | --- | --- | --- | --- | --- |
|  |  |  |  |  | Con | CS | Rie |  |  |  |  |  | Con | CS | Rie |  |
| Start | -6 | 6 | 6 | 66 | 31 | 23 | 2 | 6 | 2 | 2 | -2.3 | 12 | 1.9 | 1.3 | 1.7 | 3.5 |
| Stop | 0 | 14 | 11 | 66 | 31 | 23 | 7 | 6 | 7 | 5 | -1.7 | 24 | 2.4 | 1.8 | 2.2 | 5.0 |
| Step | 2 | 2 | 1 | 0 | 0 |  |  | 0 | 1 | 1 | 0.1 | 2 | 0.1 |  |  | 0.5 |
| n | 4 | 5 | 6 | 1 | 1 |  |  | 1 | 6 | 4 | 7 | 7 | 6 |  |  | 4 |
| Equation | Eq. 2 |  | Eqs. 2, 7 |  | Eq. 3 |  |  | Eqs. 1, 3, 4, 5 | Eq. 3 |  | Eqs. 4, 5 |  | Eq. 6 |  |  |  |

## 3. Results

The NYUS.1 model proposed here using optimized parameters for each cultivar based on calibration dataset (**Table 2**, **Fig. 1**) produced good predictions of cold hardiness for the validation dataset of the three cultivars studied (RMSE_val_ ranging from 2.12 °C to 2.72 °C, bias from –1.46 °C to 1.28 °C; **Table 2**, **Figs. 3 and 4A**).

**Table 2.** Parameter value combinations that minimize root mean square error for three grapevine cultivars in calibration datasets. Years used in calibration and validation datasets.

| Cultivar | $T_{th_{low}}$ | $T_{th_{high}}$ | $b$ | $c$ | $CH_{abs\ max}$ | $CH_{bud\ set}$ | $d$ | $f$ | $g$ | $h$ | $k_{deacc}$ | $a$ | Calibration | Validation |
| --- | --- | --- | --- | --- | --- | --- | --- | --- | --- | --- | --- | --- | --- | --- |
| ‘Concord’ | 0 | 12 | 8 | 66 | −31 | −6 | 3 | 5 | −2.1 | 24 | 1.9 | 3.5 | 2012-2013,<br>2013-2014,<br>2016-2017,<br>2017-2018,<br>2019-2020 | 2014-2015,<br>2018-2019,<br>2020-2021 |
| ‘Cabernet<br>Sauvignon’ | 0 | 8 | 7 | 66 | −23 | −6 | 6 | 3 | −2.2 | 22 | 1.8 | 5.0 | 2016-2017,<br>2017-2018,<br>2018-2019,<br>2020-2021 | 2015-2016,<br>2019-2020 |
| ‘Riesling’ | 0 | 12 | 7 | 66 | −26 | −6 | 3 | 4 | −1.9 | 24 | 2.2 | 5.0 | 2012-2013,<br>2014-2015,<br>2017-2018,<br>2018-2019,<br>2019-2020,<br>2020-2021 | 2013-2014,<br>2015-2016,<br>2016-2017 |

**Figure 3.**
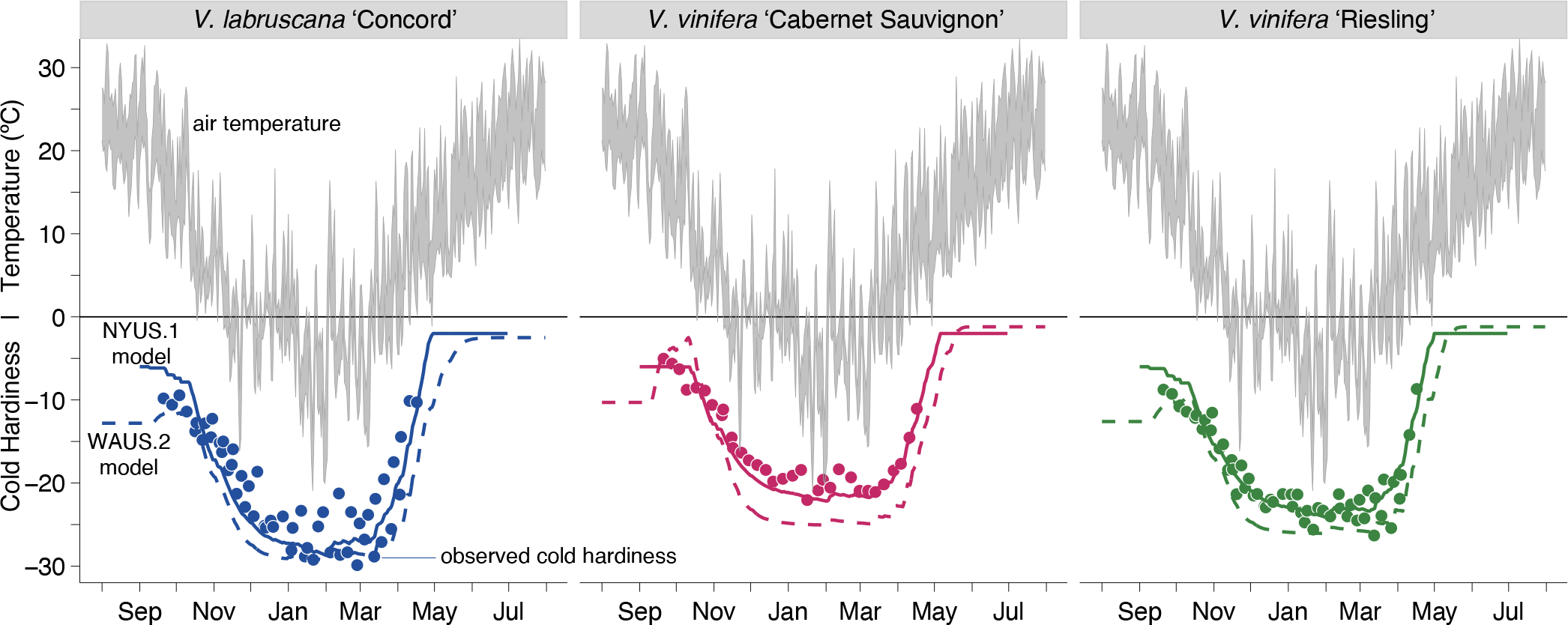
Observed cold hardiness, based on average low temperature exotherms (points), plotted with predicted bud cold hardiness according to NYUS.1 model (solid lines) and WAUS.2 model (dashed lines) with daily maximum and minimum temperature range (grey ribbons) in 2018-2019 for three grapevine cultivars.

**Figure 4.**
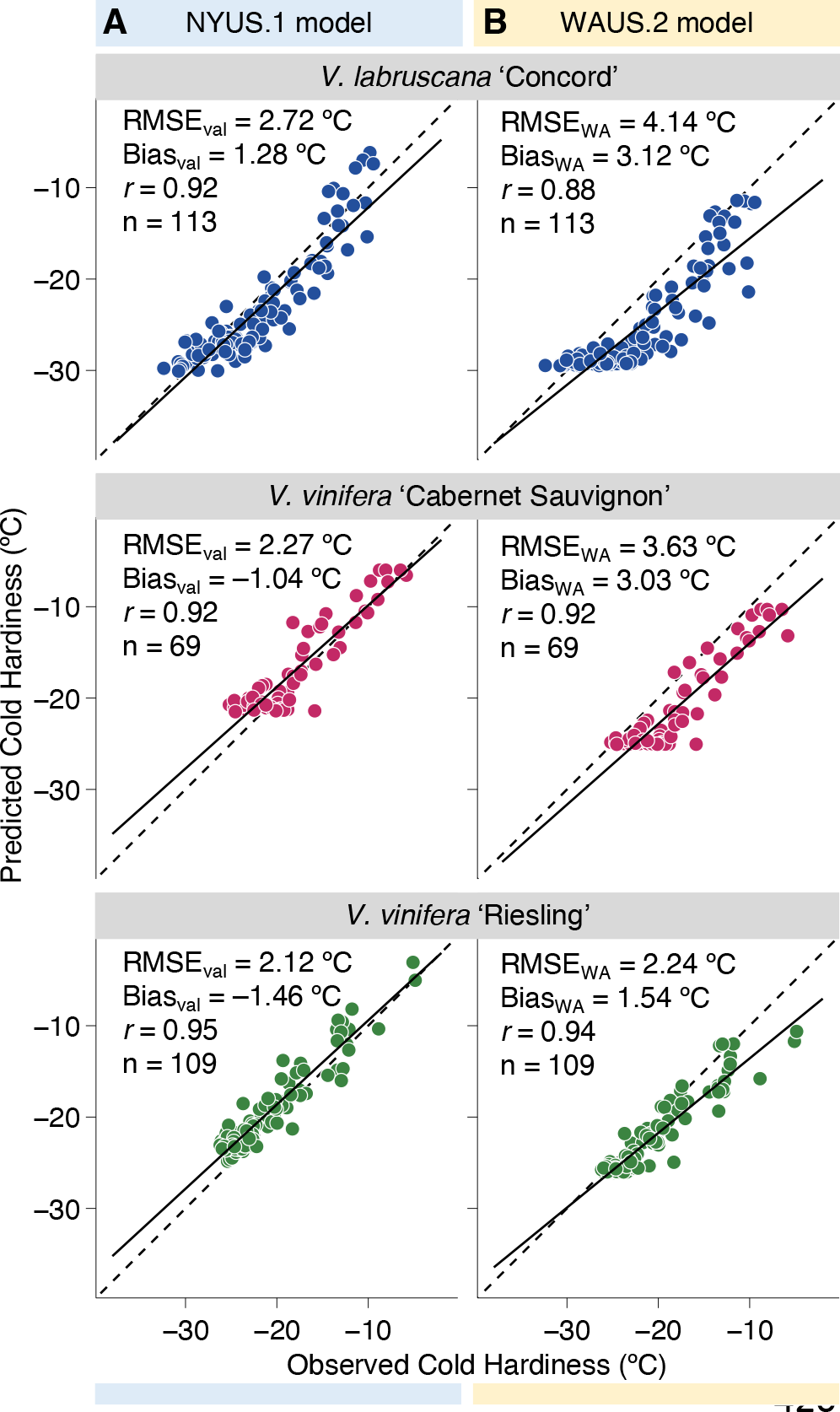
Comparison between predicted and observed bud cold hardiness, reported as average low temperature exotherm (LT_50_), for three grapevine cultivars using both the (**A**) NYUS.1 model and (**B**) WAUS.2 model. Statistics are listed in each panel, including root mean square error (RMSE), bias, correlation coefficient (r), and sample size (n). The prediction and observation correlations are illustrated (solid black line) along with a one-to-one reference (dashed black line).

The values for RMSE_cal_ (2.09 °C, 1.47 °C, 1.61 °C for ‘Concord’, ‘Cabernet Sauvignon’, and ‘Riesling’, respectively) and RMSE_val_ (**Fig. 4A**) were similar within each cultivar. This demonstrates the model and parameters chosen are robust, thus predicting similar variability in both datasets, dealing well with the year-to-year weather variations. In absolute values, predictions produced were slightly better for the two *V. vinifera* cultivars compared to ‘Concord’. This may be a result of the higher responsiveness of ‘Concord’ compared to either *V. vinifera* (greater changes in *CH* over time), as well as its overall greater cold hardiness.

Compared to the existing model, WAUS.2, the newly developed NYUS.1 model produced better predictions for ‘Concord’ and ‘Cabernet Sauvignon’, and slightly better or comparable predictions for ‘Riesling’ for data from Geneva, NY (**Figs. 3,4**). RMSE_WA_ was only computed within the validation datasets, and had values of 4.14 °C, 3.63 °C, and 2.24 °C for ‘Concord’, ‘Cabernet Sauvignon’, and ‘Riesling’, respectively.

The WAUS.2 model consistently overpredicted cold hardiness (Bias_WA_ of 3.12 °C, 3.03 °C, and 1.54 °C for ‘Concord’, ‘Cabernet Sauvignon’, and ‘Riesling’, respectively; **Fig. 4B**). This is likely due to the development of the model using data from a region with relatively warmer and drier winters. The bias may be enhanced by the WAUS.2 model’s design, where any temperature below the temperature threshold that separates acclimation and deacclimation can lead to full acclimation to the maximum cold hardiness. In the present model, low temperatures have both a different maximum cold hardiness possible (**Eqs. 2 and 3**) and different acclimation rates (due to the effect of maximum cold hardiness on **Eqs. 4 and 5**). Our model, although selected based on lowest RMSE_cal_, also had low bias for the three cultivars.

In the same season, ‘Concord’ gains more cold hardiness, and does so faster, than the two *V. vinifera* despite the same starting cold hardiness. This effect is observed in the generally lower *CH_max_* (**Fig. 1B**) and higher rates of acclimation in response to *ΔT_acc_* (**Fig. 1D**). ‘Concord’ also loses its cold hardiness faster than the *V. vinifera*, while ‘Riesling’ is also faster than ‘Cab. Sauvignon’. Within the optimized parameters, ‘Riesling’ has a higher 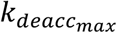 than ‘Concord’. However, because of the higher *a* parameter for **Eq. 6** (**Table 2**), ‘Riesling’ has lower deacclimation rates at temperatures between 0 °C and 17.5 °C (**Fig. 1E**), which are temperatures that largely predominate during the spring, when the majority of deacclimation occurs.

While all combinations of parameters were tested for optimization within each cultivar, some parameters were found to be very similar between the cultivars for the NYUS.1 model. This could suggest that the model is realistic in describing a physiological process that is conserved among the different cultivars within a species (‘Cabernet Sauvignon’ and ‘Riesling’), and among grapevine species (here *V. vinifera* and *V. labruscana*). ‘Cabernet Sauvignon’ had a lower 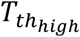 than ‘Concord’ and ‘Riesling’, but the rate of change in 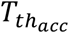 in response to chilling (*b*) was very similar. The parameter *b*, which defines the slope of the log-logistic associated here with both 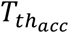 and *Ψ_deacc_*, has previously been suggested to be the same for all species for deacclimation (Kovaleski, 2022), leaving regulation of magnitudes of deacclimation (effective deacclimation) to the effect of temperature alone on the rate of deacclimation, *k_deacc_*.

## 4. Discussion

Accurate modeling of cold hardiness is an important step in understanding adaptation of species and cultivars in future climates (Wolkovich et al., 2018; Körner, 2021). However, modeling of this trait is also helpful for growers in guiding management practices and in cultivar choice for different regions in current climates. Here we used known deacclimation responses (Kovaleski et al., 2018; Kovaleski, 2022; North et al., 2022) and posited acclimation responses, along with a medium-term dataset with three cultivars that have relatively different cold hardiness behaviors to create a new cold hardiness prediction model for grapevine buds.

Our new model (NYUS.1 model) outperforms the currently available model for grapevine cold hardiness (WAUS.2; Ferguson et al., 2014) using New York state data. The WAUS.2 model consistently overestimates bud cold hardiness in New York, which consequently can underestimate freeze injury risk and occurrence. An overestimated cold hardiness could mislead adaptation of production systems (e.g. site selection or cultivar evaluation) and management decisions (e.g. spring pruning decisions). This is especially consequential as extreme and erratic weather events during winter become more frequent under changing climates (Casson et al., 2019; IPCC, 2022).

Parameters here are optimized using field data, as in other modeling studies (Ferguson et al., 2011; Ferguson et al., 2014; North et al., 2021). This was done despite some data available on experimentally determined deacclimation rates for the two *V. vinifera* cultivars (Kovaleski et al., 2018) and a few hybrids (North et al., 2022), though not ‘Concord’. However, the estimated deacclimation rates based on **Eq. 6** for either *V. vinifera* cultivars were comparable to values previously reported (Kovaleski et al., 2018). This indicates field data, associated with models that prioritize the realism model dimension, can estimate parameters equivalent to controlled experiments.

The modulation of acclimation rate based on a logistic regression in response to time has a similar effect here as that of the Theta (*θ*) term in the WAUS.2 model, effectively attenuating the approach to any given cold hardiness. However, because each temperature elicits a different cold hardiness in our model (**Eqs. 2 and 3**), this effect is dynamic and can occur at any cold hardiness and any temperature, rather than just the asymptotical approach to absolute maximum and minimum cold hardiness as in Ferguson et al. (2014). This may improve the generalizability of our model in applications to regions other than New York. Deployment of the model to various cold hardiness monitoring programs (e.g., MI, PA, OH, CO, and Canada) for validation of fit would enable validation of these assumptions. Given that recent studies (Kovaleski et al., 2018; Kovaleski, 2022; North et al., 2022; North and Kovaleski, 2022) have demonstrated that cold hardiness status (as measured with DTA) is lost prior to budbreak, further development in cold hardiness models [e.g., adding a cold hardiness where budbreak is observed (Kovaleski et al., 2022)] could lead to tests using budbreak phenology datasets in regions where cold hardiness is not monitored as well.

The optimized 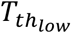 was 0 °C for all cultivars, which is the upper limit established. For a best fit of the model, it would be ideal to increase the range for iterations such that RMSE may be further minimized, ideally finding a global minimum. However, a higher 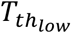 would mean that the small deacclimations that happen at low but positive temperatures might have been counterbalanced by the effect of acclimation. By limiting 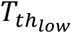 to 0 °C we prevented this effect, which would otherwise be particularly consequential during the spring when plants are fully chilled. Should data be gathered in warmer regions with lower seasonal chill accumulation, this limit should be evaluated and possibly be re-parameterized to remove these effects.

The calibration and validation datasets were not the same for all cultivars (**Table 2**; **Supplementary Figs. 1-3**). This was due to data not having been collected the same number of years for all cultivars. However, enough variation in the years available for the three cultivars appear to allow for good parameterization of the models. For example, within the calibration dataset for ‘Concord’, 2012-2013 is a much milder winter both in minimum temperature and the length of exposure to very low temperatures as compared to 2013-2014 (**Supplementary Fig. 1**).

Here, the transition of endodormancy to ecodormancy occurs quantitatively through the accumulation of chill that modulates the rates of deacclimation (*k_deacc_*) through the deacclimation potential (*Ψ_deacc_*). This is very different from the WAUS.1 and WAUS.2 models, where a qualitative transition occurs. It is likely that a qualitative transition works well as an assumption within a single region, given some regularity in the level of chill accumulation across years. However, for a single model to accomplish the estimation of deacclimation rates in regions of lower chill accumulation and higher chill accumulation, a quantitative transition is required based on experimental results (Kovaleski et al., 2018; Kovaleski, 2022, North et al., 2022). Field data from different regions, particularly those where low chilling accumulation occur will be required to test this. It is also important to note that some management practices that “supplement chilling” and break dormancy (e.g., application of hydrogen cyanamide) have not been studied from a cold hardiness context, and therefore we do not expect the model to excel if applied to regions where such treatments occur until effects of dormancy breaking treatments has been studied.

The dynamic model was used here to estimate chilling accumulation, due to its reported higher transferability to warmer environments (Luedeling and Brown, 2011). Within high chill accumulation regions, it has been reported to be highly correlated to other chilling models, such as “Utah” (Richardson et al., 1974) and “North Carolina” (Shaltout and Unrath, 1983) [e.g., in Boston, MA, USA (Kovaleski, 2022), and Madison, WI, USA (North et al., 2022)]. However, it is still possible that all chill models are incorrectly describing chilling accumulation, and therefore this may be a source of error for predictions, especially in regions with low chill accumulation. Deacclimation experiments, such as in other studies by the present authors (Kovaleski et al., 2018; Kovaleski, 2022; North et al., 2022), that use material collected from warmer climates should help elucidate the dynamics of chilling. Should the chilling accumulation models change, parameter changes within the model presented here may be required.

In our calibration dataset, the rounded average of the 15 most cold hardy points for ‘Concord’ is −31 °C, and the lowest observed cold hardiness was −31.4 °C. This is below the maximum possible using the WAUS.2 model for that cultivar (−29.5 °C). This suggests the maximum cold hardiness for this genotype may not be achieved in Washington’s conditions, which are relatively warmer compared to upstate New York. The higher value of cold hardiness being the maximum possible in their model results in a hard limit seen in the lower end of the observed vs. predicted plots for the WAUS.2 model (**Fig. 4B**). The opposite is observed in ‘Cabernet Sauvignon’, however, where the present model uses a maximum cold hardiness of −23 °C (although this value would be −26 °C if estimated from the validation dataset), whereas the Washington model uses a value of −25.1 °C. In the currently proposed model, besides having a lower maximum, this is not a hard limit given the possibility of a low acclimation rate at any temperature.

The values estimated for the slope associated with the log logistic for chill responses were similar to those previously reported (Kovaleski, 2022). Here, both acclimation and deacclimation were guided by the same slope, providing the phased integration of both, which may have led to slight differences from measured effects. Future studies in acclimation could help understand whether the response should be maintained as varying alongside deacclimation or not.

The similarity in parameter values across cultivars for acclimation and deacclimation, particularly the portions that *do not describe magnitude* (i.e., slope for 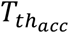, slope and intercept for *CH^*^*, and slope for *Ψ_deacc_*), suggests a conserved physiology if the present model (NYUS.1) is correctly approximating responses. This has been suggested for the dormancy and deacclimation across the land plant phylogeny (Kovaleski, 2022), though without using multiple genotypes within a species. Deacclimation appears to be a simpler process, where no temperature sensing is required, only temperature perception (Penfield, 2008). It remains to be tested whether acclimation has conserved mechanisms that can easily translate to mathematical models of the process, though acclimation appears to be more difficult to describe and elicit in controlled environment experiments.

Variability in model performance across regions is at least partially a consequence of model development with datasets from a single region. However, there is a lack of freely available datasets of cold hardiness from different climates that restricts improvements in model development, unlike with other types of data such as spring phenology (e.g., Pan European Phenology Project and USA’s National Phenology Network). Therefore, we are making our dataset public so other efforts may use this medium-term dataset (https://github.com/apkovaleski/NYUS.1_CHmodel; Kovaleski et al., 2022). It is also noticeable that most cold hardiness datasets are produced in higher latitudes and temperate climates where considerably low temperatures occur during winter [e.g., Washington (Ferguson et al., 2014), Wisconsin (North et al., 2021, 2022) and New York states within the US; Chile (Rubio and Pérez, 2020)]. There is a need for new data to be collected from warmer locations in lower latitudes such that parameters can be optimized based on warmer subtropical (or even tropical) climates. It is possible that the model proposed here, the existing WAUS.2 model (Ferguson et al., 2014), or regional adaptations (North et al., 2021) perform variably across different regions, and therefore models should continue to be used in conjunction in new regions to validate and tune predictions. As models continue to be improved, convergence of responses should result in high confidence of predictions.

Our model uses only air temperature-related parameters to describe cold hardiness. Some variation in cold hardiness can also be explained by temperature mediated aspects of microclimate and daylength, such as through radiative warming (Grace, 2006; Quiñones et al., 2019; Vitasse et al., 2021), wind (Grace, 2006), water availability, and daily amplitude in temperature (Gonzalez Antivilo et al., 2017; Matsubara 2018). In addition to direct effects on cold hardiness, such factors may indirectly affect cold hardiness through effects on dormancy. However, integrating these relatively minor factors influencing cold hardiness changes into models, while reducing prediction error somewhat, also increase a model’s computational complexity and potentially make the results of parameter computing less generalizable.

## 5. Conclusions

The NYUS.1 model produced here demonstrates a robust manner of predicting cold hardiness for buds of three different grapevine cultivars. This model integrates novel deacclimation concepts discussed in recent literature (Kovaleski et al., 2018; Kovaleski, 2022; North et al., 2022). It is possible that future research regarding acclimation could lead to a better understanding of the processes leading to gains in cold hardiness and subsequently improved modeling approaches. However, acclimation seems to be an elusive process to date and coincides with other significant physiological changes that are only partially understood (e.g., dormancy induction). Therefore, further research in acclimation and cold hardiness modeling will need to continually assess the balance between model realism and model accuracy. The NYUS.1 model proposed here provides accurate cold hardiness predictions in the meantime and a framework on which to build. Better predictions were produced using the NYUS.1 model when compared to the WAUS.2 model (Ferguson et al., 2014). Thus, the proposed model may be used in lieu of, or in addition to, existing models to aid in decision making steps. Despite having been developed using regional data, the selection of similar parameters across cultivars could indicate a representative description of physiological processes that may also promote transferability into other regions with some caveats, such as errors in chilling accumulation models.

## Acknowledgements

We would like to thank Hanna Martens, Bill Wilsey, Lex Pike, and Kathleen Deys for help collecting and processing grape bud tissue for field cold hardiness measurements. We would also like to thank Ravines Vineyard in Geneva, NY for access to *V. vinifera* ‘Cabernet Sauvignon’ and ‘Riesling’ vines. This work was partially supported by the Office of the Vice Chancellor for Research and Graduate Education at the University of Wisconsin–Madison with funding from the Wisconsin Alumni Research Foundation, and through the USDA ARS appropriated project 1910-21220-006-00D.

## Supplementary Figure Legends

**Supplementary Figure 1**. *Vitis labruscana* ‘Concord’ validation, calibration, and N/A datasets. Observed cold hardiness, based on average low temperature exotherms (blue points) plotted with predicted bud cold hardiness according to NYUS.1 model (solid blue lines) and WAUS.2 model (dashed blue lines) with daily maximum and minimum temperature range (grey ribbons).

**Supplementary Figure 2**. *Vitis vinifera* ‘Cabernet Sauvignon’ validation, calibration, and N/A datasets. Observed cold hardiness, based on average low temperature exotherms (magenta points) plotted with predicted bud cold hardiness according to NYUS.1 model (solid magenta lines) and WAUS.2 model (dashed magenta lines) with daily maximum and minimum temperature range (grey ribbons).

**Supplementary Figure 3**. *Vitis vinifera* ‘Riesling’ validation, calibration, and N/A datasets. Observed cold hardiness, based on average low temperature exotherms (green points) plotted with predicted bud cold hardiness according to NYUS.1 model (solid green lines) and WAUS.2 model (dashed green lines) with daily maximum and minimum temperature range (grey ribbons).

