## Supplementary Figures 1-3 for "Development of a new cold hardiness prediction model for grapevine using phased integration of acclimation and deacclimation responses"

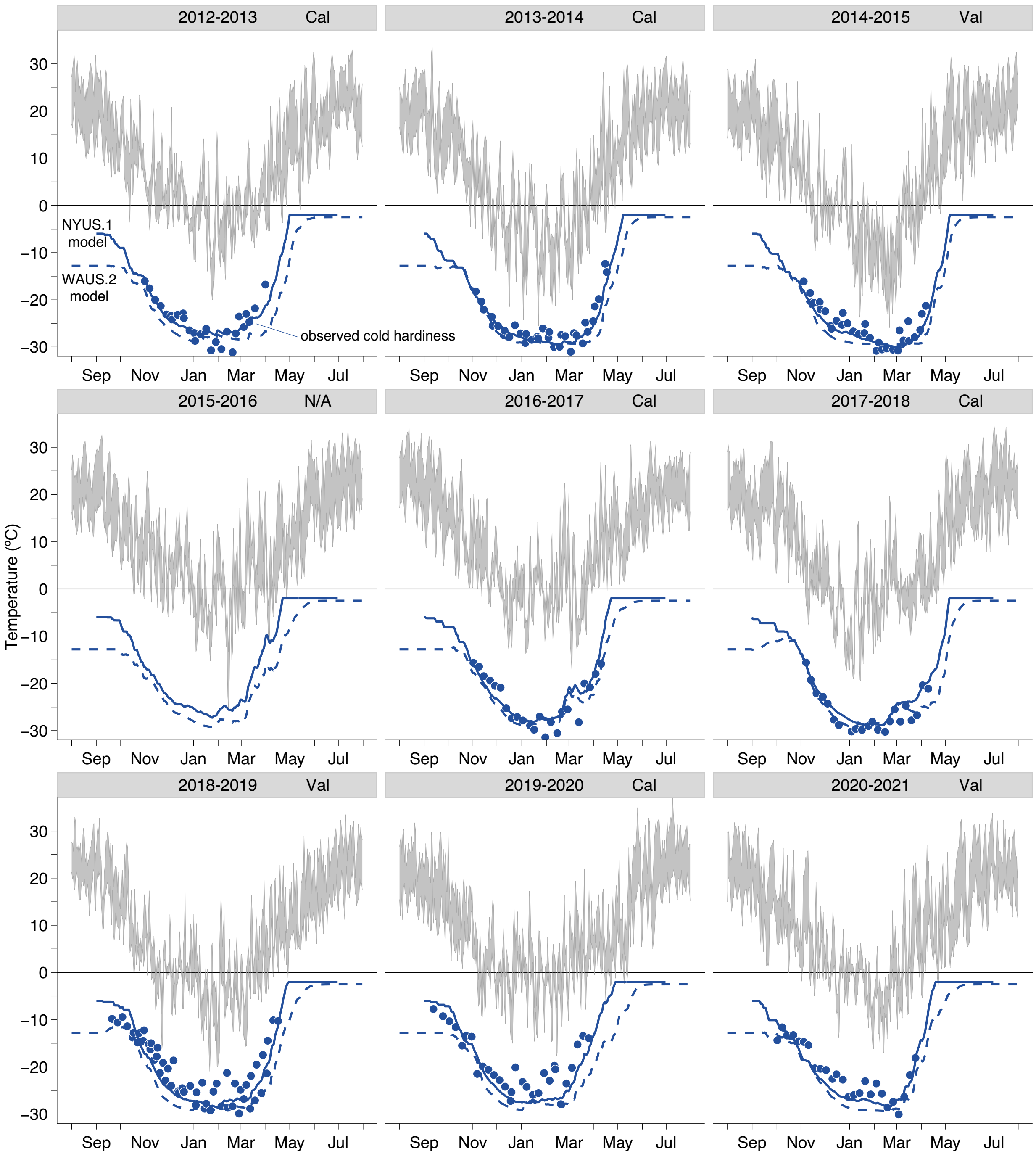


**Supplementary Figure 1.** *Vitis labruscana* 'Concord' validation, calibration, and N/A datasets. Observed cold hardiness, based on average low temperature exotherms (blue points) plotted with predicted bud cold hardiness according to NYUS.1 model (solid blue lines) and WAUS.2 model (dashed blue lines) with daily maximum and minimum temperature range (grey ribbons).

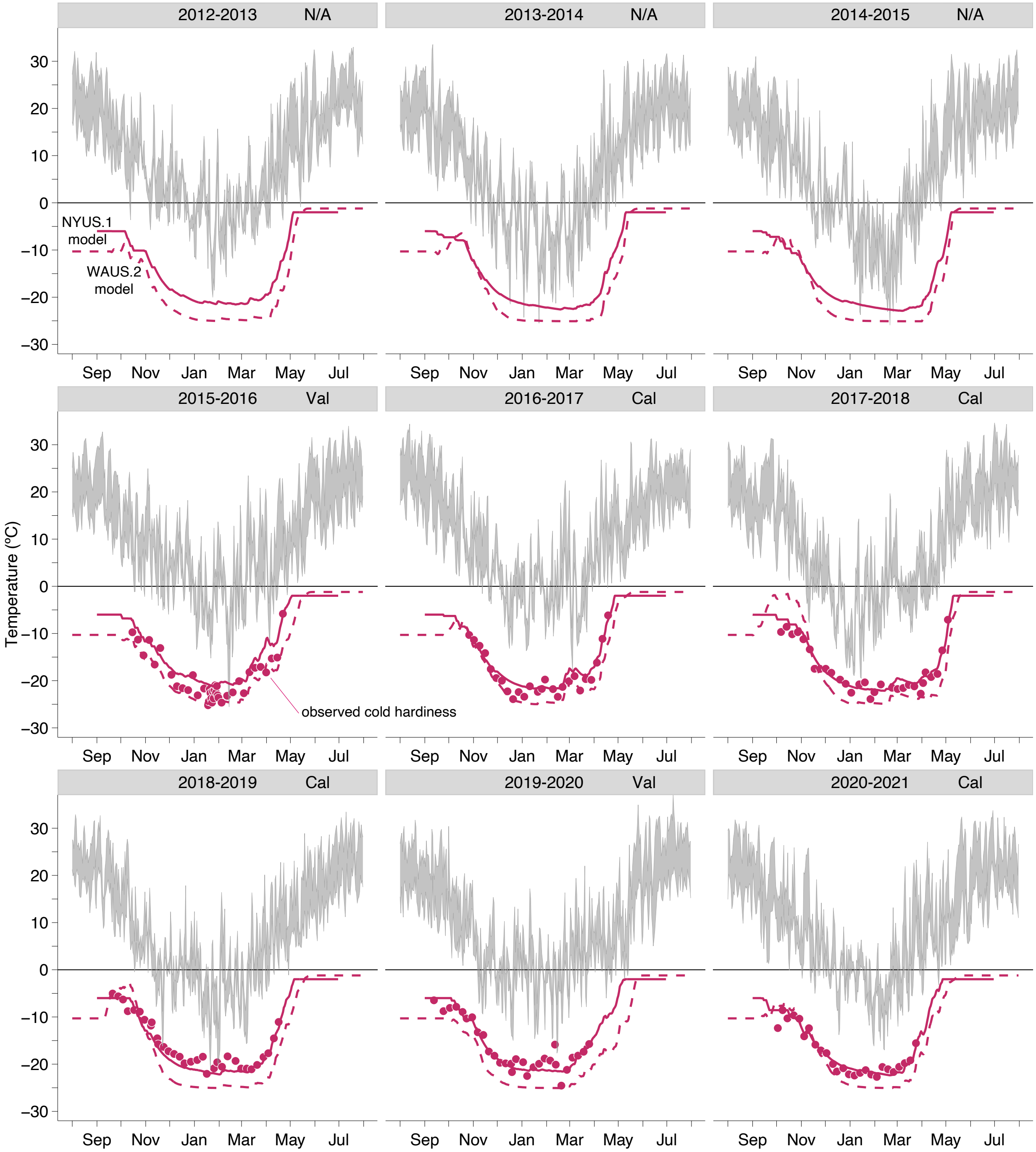


**Supplementary Figure 2.** *Vitis vinifera* 'Cabernet Sauvignon' validation, calibration, and N/A datasets. Observed cold hardiness, based on average low temperature exotherms (magenta points) plotted with predicted bud cold hardiness according to NYUS.1 model (solid magenta lines) and WAUS.2 model (dashed magenta lines) with daily maximum and minimum temperature range (grey ribbons).

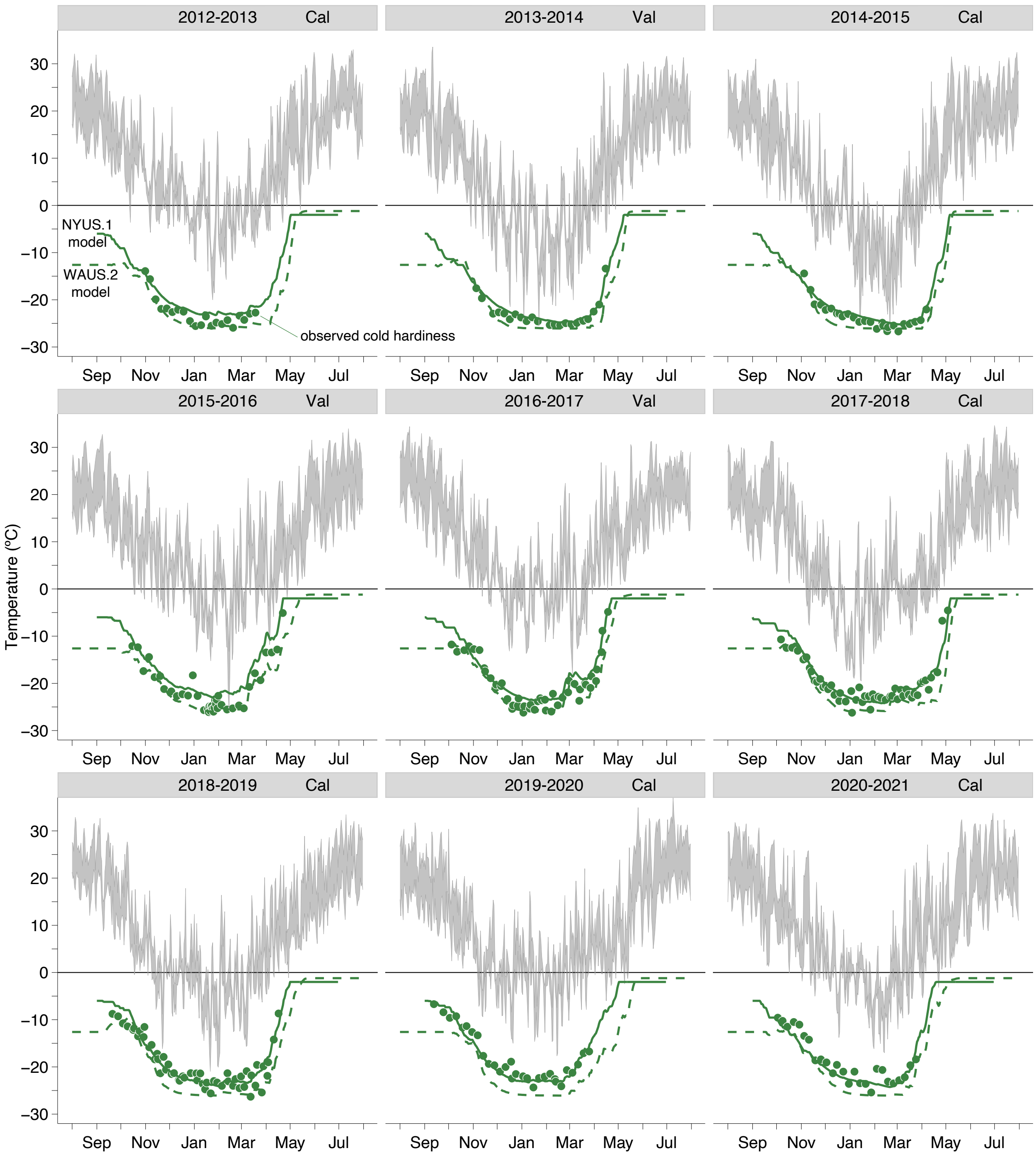


**Supplementary Figure 3.** *Vitis vinifera* 'Riesling' validation, calibration, and N/A datasets. Observed cold hardiness, based on average low temperature exotherms (green points) plotted with predicted bud cold hardiness according to NYUS.1 model (solid green lines) and WAUS.2 model (dashed green lines) with daily maximum and minimum temperature range (grey ribbons).
