## Supplementary Table 1 for "Development of a new cold hardiness prediction model for grapevine using phased integration of acclimation and deacclimation responses"

**Supplementary Table 1.** Mean, standard deviation, and n of observed cold hardiness (LTE) by cultivar, date, and program.

| Cultivar | Date | Dataset | Program | LTE | Standard deviation | n |
|---|---|---|---|---|---|---|
| *V. labruscana* 'Concord' | 10/31/12 | 2012-2013 | Martinson | -16.05 | 0.87 | 30 |
| | 11/6/12 | 2012-2013 | Londo | -17.80 | 1.61 | 8 |
| | 11/6/12 | 2012-2013 | Martinson | -17.52 | 1.24 | 30 |
| | 11/13/12 | 2012-2013 | Martinson | -20.01 | 2.01 | 30 |
| | 11/20/12 | 2012-2013 | Londo | -18.04 | 2.83 | 9 |
| | 11/20/12 | 2012-2013 | Martinson | -22.30 | 1.27 | 30 |
| | 11/27/12 | 2012-2013 | Martinson | -23.11 | 1.32 | 30 |
| | 12/3/12 | 2012-2013 | Londo | -23.42 | 0.43 | 8 |
| | 12/4/12 | 2012-2013 | Martinson | -24.17 | 1.15 | 30 |
| | 12/11/12 | 2012-2013 | Martinson | -23.20 | 1.84 | 30 |
| | 12/18/12 | 2012-2013 | Martinson | -22.88 | 1.34 | 30 |
| | 12/19/12 | 2012-2013 | Londo | -23.90 | 0.63 | 2 |
| | 12/26/12 | 2012-2013 | Martinson | -26.48 | 0.94 | 30 |
| | 1/1/13 | 2012-2013 | Londo | -27.04 | 1.28 | 2 |
| | 1/2/13 | 2012-2013 | Martinson | -28.70 | 1.38 | 30 |
| | 1/9/13 | 2012-2013 | Martinson | -27.29 | 1.35 | 30 |
| | 1/15/13 | 2012-2013 | Martinson | -26.68 | 1.39 | 30 |
| | 1/16/13 | 2012-2013 | Londo | -26.13 | 1.62 | 9 |
| | 1/22/13 | 2012-2013 | Martinson | -30.72 | 2.02 | 30 |
| | 1/28/13 | 2012-2013 | Martinson | -28.96 | 1.82 | 30 |
| | 2/4/13 | 2012-2013 | Londo | -28.51 | 1.66 | 9 |
| | 2/4/13 | 2012-2013 | Martinson | -31.33 | 1.89 | 20 |
| | 2/11/13 | 2012-2013 | Martinson | -26.77 | 1.40 | 30 |
| | 2/18/13 | 2012-2013 | Martinson | -31.13 | 1.19 | 30 |
| | 2/22/13 | 2012-2013 | Londo | -27.13 | 1.86 | 9 |
| | 2/26/13 | 2012-2013 | Martinson | -23.55 | 1.64 | 30 |
| | 3/4/13 | 2012-2013 | Martinson | -25.78 | 1.34 | 30 |
| | 3/7/13 | 2012-2013 | Londo | -22.98 | 1.55 | 8 |
| | 3/11/13 | 2012-2013 | Martinson | -24.69 | 2.45 | 30 |
| | 3/18/13 | 2012-2013 | Martinson | -21.81 | 1.74 | 30 |
| | 3/31/13 | 2012-2013 | Londo | -16.78 | 3.32 | 8 |
| | 11/4/13 | 2013-2014 | Martinson | -18.23 | 1.40 | 27 |
| | 11/11/13 | 2013-2014 | Martinson | -20.42 | 1.25 | 27 |
| | 11/14/13 | 2013-2014 | Londo | -22.08 | 1.21 | 10 |
| | 11/24/13 | 2013-2014 | Londo | -23.61 | 1.12 | 10 |
| | 11/25/13 | 2013-2014 | Martinson | -25.49 | 1.41 | 27 |

| | | | | | |
|---|---|---|---|---|---|
| 12/2/13 | 2013-2014 | Martinson | -25.59 | 2.05 | 27 |
| 12/9/13 | 2013-2014 | Martinson | -27.51 | 1.36 | 27 |
| 12/10/13 | 2013-2014 | Londo | -26.53 | 0.79 | 7 |
| 12/16/13 | 2013-2014 | Martinson | -27.89 | 2.06 | 27 |
| 12/23/13 | 2013-2014 | Londo | -24.99 | 0.84 | 7 |
| 12/23/13 | 2013-2014 | Martinson | -25.51 | 1.82 | 27 |
| 12/30/13 | 2013-2014 | Martinson | -27.12 | 1.33 | 27 |
| 1/5/14 | 2013-2014 | Londo | -29.15 | 0.86 | 9 |
| 1/6/14 | 2013-2014 | Martinson | -27.24 | 2.38 | 27 |
| 1/13/14 | 2013-2014 | Martinson | -28.48 | 1.83 | 27 |
| 1/20/14 | 2013-2014 | Martinson | -27.86 | 1.11 | 27 |
| 1/21/14 | 2013-2014 | Londo | -28.15 | 1.16 | 5 |
| 1/27/14 | 2013-2014 | Martinson | -26.07 | 2.54 | 27 |
| 2/3/14 | 2013-2014 | Martinson | -28.03 | 1.24 | 27 |
| 2/4/14 | 2013-2014 | Londo | -26.81 | 1.53 | 9 |
| 2/10/14 | 2013-2014 | Martinson | -30.03 | 1.66 | 27 |
| 2/17/14 | 2013-2014 | Martinson | -29.97 | 2.71 | 27 |
| 2/19/14 | 2013-2014 | Londo | -27.44 | 0.80 | 7 |
| 2/24/14 | 2013-2014 | Martinson | -27.71 | 1.87 | 27 |
| 3/3/14 | 2013-2014 | Martinson | -31.02 | 2.41 | 22 |
| 3/7/14 | 2013-2014 | Londo | -27.07 | 0.87 | 7 |
| 3/10/14 | 2013-2014 | Martinson | -27.51 | 1.44 | 27 |
| 3/18/14 | 2013-2014 | Martinson | -29.24 | 1.81 | 25 |
| 3/21/14 | 2013-2014 | Londo | -24.83 | 1.10 | 10 |
| 3/24/14 | 2013-2014 | Martinson | -26.81 | 1.77 | 27 |
| 3/31/14 | 2013-2014 | Martinson | -24.54 | 1.25 | 27 |
| 4/2/14 | 2013-2014 | Londo | -21.41 | 1.91 | 8 |
| 4/7/14 | 2013-2014 | Martinson | -19.85 | 1.47 | 27 |
| 4/15/14 | 2013-2014 | Martinson | -12.38 | 3.16 | 27 |
| 4/17/14 | 2013-2014 | Londo | -14.14 | 0.56 | 7 |
| 11/4/14 | 2014-2015 | Martinson | -16.13 | 1.20 | 30 |
| 11/12/14 | 2014-2015 | Martinson | -18.56 | 1.87 | 30 |
| 11/14/14 | 2014-2015 | Londo | -20.57 | 2.52 | 9 |
| 11/17/14 | 2014-2015 | Martinson | -20.64 | 2.53 | 30 |
| 11/24/14 | 2014-2015 | Londo | -20.51 | 2.22 | 8 |
| 11/25/14 | 2014-2015 | Martinson | -22.02 | 2.35 | 30 |
| 12/1/14 | 2014-2015 | Martinson | -22.42 | 2.43 | 30 |
| 12/8/14 | 2014-2015 | Martinson | -25.53 | 2.09 | 30 |
| 12/9/14 | 2014-2015 | Londo | -26.13 | 1.37 | 6 |
| 12/15/14 | 2014-2015 | Martinson | -24.44 | 2.54 | 30 |

| | | | | | |
|---|---|---|---|---|---|
| 12/22/14 | 2014-2015 | Martinson | -25.31 | 1.73 | 30 |
| 12/23/14 | 2014-2015 | Londo | -22.77 | 1.75 | 10 |
| 12/29/14 | 2014-2015 | Martinson | -25.00 | 2.46 | 30 |
| 1/5/15 | 2014-2015 | Londo | -26.26 | 0.83 | 6 |
| 1/5/15 | 2014-2015 | Martinson | -26.80 | 1.84 | 30 |
| 1/12/15 | 2014-2015 | Martinson | -27.27 | 2.31 | 20 |
| 1/19/15 | 2014-2015 | Martinson | -27.05 | 1.92 | 30 |
| 1/20/15 | 2014-2015 | Londo | -25.15 | 2.96 | 9 |
| 1/26/15 | 2014-2015 | Martinson | -28.04 | 2.75 | 30 |
| 2/3/15 | 2014-2015 | Martinson | -30.80 | 2.37 | 30 |
| 2/6/15 | 2014-2015 | Londo | -29.08 | 2.73 | 9 |
| 2/10/15 | 2014-2015 | Martinson | -30.52 | 2.25 | 30 |
| 2/16/15 | 2014-2015 | Martinson | -30.31 | 3.19 | 29 |
| 2/21/15 | 2014-2015 | Londo | -32.38 | 1.18 | 3 |
| 2/24/15 | 2014-2015 | Martinson | -30.59 | 3.63 | 30 |
| 3/2/15 | 2014-2015 | Martinson | -30.76 | 2.41 | 30 |
| 3/4/15 | 2014-2015 | Londo | -26.48 | 2.77 | 10 |
| 3/9/15 | 2014-2015 | Martinson | -28.61 | 1.79 | 30 |
| 3/15/15 | 2014-2015 | Londo | -24.53 | 1.58 | 10 |
| 3/16/15 | 2014-2015 | Martinson | -28.72 | 2.16 | 30 |
| 3/23/15 | 2014-2015 | Martinson | -27.92 | 3.07 | 30 |
| 3/30/15 | 2014-2015 | Martinson | -26.40 | 2.34 | 30 |
| 4/1/15 | 2014-2015 | Londo | -22.96 | 2.18 | 11 |
| 4/6/15 | 2014-2015 | Martinson | -21.28 | 2.73 | 30 |
| 11/1/16 | 2016-2017 | Martinson | -15.66 | 1.45 | 30 |
| 11/8/16 | 2016-2017 | Martinson | -16.45 | 2.52 | 30 |
| 11/14/16 | 2016-2017 | Martinson | -18.49 | 1.28 | 30 |
| 11/22/16 | 2016-2017 | Martinson | -19.43 | 1.91 | 30 |
| 11/28/16 | 2016-2017 | Martinson | -20.52 | 1.55 | 30 |
| 12/5/16 | 2016-2017 | Martinson | -20.89 | 2.09 | 30 |
| 12/12/16 | 2016-2017 | Martinson | -25.24 | 1.79 | 30 |
| 12/19/16 | 2016-2017 | Martinson | -27.40 | 2.03 | 30 |
| 12/26/16 | 2016-2017 | Martinson | -27.12 | 2.40 | 30 |
| 1/2/17 | 2016-2017 | Martinson | -27.86 | 4.03 | 30 |
| 1/11/17 | 2016-2017 | Martinson | -28.94 | 1.72 | 30 |
| 1/16/17 | 2016-2017 | Martinson | -29.82 | 1.75 | 30 |
| 1/23/17 | 2016-2017 | Martinson | -27.03 | 1.09 | 30 |
| 1/30/17 | 2016-2017 | Martinson | -31.43 | 1.75 | 30 |
| 2/6/17 | 2016-2017 | Martinson | -28.21 | 1.43 | 30 |
| 2/14/17 | 2016-2017 | Martinson | -30.55 | 1.95 | 30 |

| | | | | | |
|---|---|---|---|---|---|
| 2/20/17 | 2016-2017 | Martinson | -26.02 | 1.84 | 30 |
| 2/27/17 | 2016-2017 | Martinson | -25.53 | 3.06 | 30 |
| 3/13/17 | 2016-2017 | Martinson | -28.25 | 3.44 | 30 |
| 3/20/17 | 2016-2017 | Martinson | -20.05 | 1.95 | 30 |
| 3/27/17 | 2016-2017 | Martinson | -20.82 | 2.72 | 30 |
| 4/3/17 | 2016-2017 | Martinson | -17.99 | 2.58 | 30 |
| 4/10/17 | 2016-2017 | Martinson | -15.86 | 3.75 | 30 |
| 11/7/17 | 2017-2018 | Martinson | -15.60 | 1.42 | 30 |
| 11/13/17 | 2017-2018 | Martinson | -19.25 | 2.15 | 30 |
| 11/20/17 | 2017-2018 | Martinson | -22.12 | 1.81 | 30 |
| 11/28/17 | 2017-2018 | Martinson | -22.86 | 2.31 | 30 |
| 12/4/17 | 2017-2018 | Martinson | -24.29 | 1.97 | 30 |
| 12/12/17 | 2017-2018 | Martinson | -27.72 | 1.93 | 30 |
| 12/18/17 | 2017-2018 | Martinson | -28.85 | 2.45 | 30 |
| 1/3/18 | 2017-2018 | Martinson | -30.20 | 2.86 | 30 |
| 1/8/18 | 2017-2018 | Martinson | -29.69 | 2.19 | 30 |
| 1/16/18 | 2017-2018 | Martinson | -29.86 | 3.14 | 30 |
| 1/24/18 | 2017-2018 | Martinson | -29.04 | 1.60 | 30 |
| 1/29/18 | 2017-2018 | Martinson | -28.14 | 1.48 | 30 |
| 2/5/18 | 2017-2018 | Martinson | -29.84 | 2.89 | 30 |
| 2/14/18 | 2017-2018 | Martinson | -30.27 | 2.78 | 30 |
| 2/20/18 | 2017-2018 | Martinson | -28.06 | 2.13 | 20 |
| 2/26/18 | 2017-2018 | Martinson | -25.53 | 2.16 | 30 |
| 3/5/18 | 2017-2018 | Martinson | -28.10 | 2.03 | 30 |
| 3/12/18 | 2017-2018 | Martinson | -24.74 | 2.07 | 30 |
| 3/19/18 | 2017-2018 | Martinson | -27.83 | 2.42 | 30 |
| 3/26/18 | 2017-2018 | Martinson | -26.77 | 2.94 | 30 |
| 4/2/18 | 2017-2018 | Martinson | -20.42 | 2.95 | 30 |
| 4/9/18 | 2017-2018 | Martinson | -21.14 | 2.68 | 30 |
| 9/20/18 | 2018-2019 | Londo | -9.82 | 2.29 | 5 |
| 9/27/18 | 2018-2019 | Londo | -10.58 | 1.90 | 5 |
| 10/3/18 | 2018-2019 | Londo | -9.44 | 2.18 | 5 |
| 10/9/18 | 2018-2019 | Londo | -11.40 | 2.12 | 5 |
| 10/16/18 | 2018-2019 | Martinson | -13.78 | 1.24 | 30 |
| 10/17/18 | 2018-2019 | Londo | -12.76 | 0.88 | 5 |
| 10/22/18 | 2018-2019 | Martinson | -14.82 | 2.34 | 30 |
| 10/24/18 | 2018-2019 | Londo | -12.82 | 0.64 | 5 |
| 10/29/18 | 2018-2019 | Martinson | -14.49 | 1.45 | 30 |
| 10/30/18 | 2018-2019 | Londo | -12.26 | 0.92 | 5 |
| 11/5/18 | 2018-2019 | Martinson | -15.19 | 2.31 | 30 |

| | | | | | |
|---|---|---|---|---|---|
| 11/7/18 | 2018-2019 | Londo | -16.28 | 1.30 | 5 |
| 11/8/18 | 2018-2019 | Londo | -15.00 | 0.52 | 5 |
| 11/12/18 | 2018-2019 | Martinson | -18.47 | 1.97 | 30 |
| 11/15/18 | 2018-2019 | Londo | -17.80 | 0.74 | 4 |
| 11/16/18 | 2018-2019 | Londo | -15.93 | 4.19 | 3 |
| 11/19/18 | 2018-2019 | Martinson | -21.29 | 1.69 | 30 |
| 11/23/18 | 2018-2019 | Londo | -19.14 | 0.91 | 5 |
| 11/26/18 | 2018-2019 | Martinson | -22.88 | 1.74 | 30 |
| 11/29/18 | 2018-2019 | Londo | -20.37 | 1.45 | 3 |
| 12/3/18 | 2018-2019 | Martinson | -23.99 | 2.18 | 30 |
| 12/6/18 | 2018-2019 | Londo | -18.64 | 3.04 | 5 |
| 12/12/18 | 2018-2019 | Martinson | -25.14 | 3.06 | 30 |
| 12/13/18 | 2018-2019 | Londo | -25.36 | 2.28 | 5 |
| 12/17/18 | 2018-2019 | Martinson | -24.52 | 1.81 | 30 |
| 12/19/18 | 2018-2019 | Londo | -25.28 | 1.91 | 10 |
| 12/27/18 | 2018-2019 | Londo | -24.02 | 1.45 | 5 |
| 1/3/19 | 2018-2019 | Martinson | -28.10 | 2.03 | 30 |
| 1/4/19 | 2018-2019 | Londo | -25.40 | 0.88 | 5 |
| 1/11/19 | 2018-2019 | Londo | -23.34 | 2.13 | 5 |
| 1/14/19 | 2018-2019 | Martinson | -28.84 | 1.89 | 30 |
| 1/16/19 | 2018-2019 | Londo | -27.81 | 0.61 | 5 |
| 1/21/19 | 2018-2019 | Martinson | -29.24 | 1.62 | 30 |
| 1/25/19 | 2018-2019 | Londo | -25.23 | 1.58 | 5 |
| 1/29/19 | 2018-2019 | Londo | -23.51 | 3.79 | 5 |
| 2/4/19 | 2018-2019 | Londo | -23.54 | 4.74 | 5 |
| 2/4/19 | 2018-2019 | Martinson | -29.16 | 1.99 | 30 |
| 2/11/19 | 2018-2019 | Londo | -21.26 | 3.77 | 10 |
| 2/12/19 | 2018-2019 | Martinson | -28.65 | 2.58 | 30 |
| 2/18/19 | 2018-2019 | Martinson | -28.34 | 3.24 | 30 |
| 2/21/19 | 2018-2019 | Londo | -23.50 | 2.91 | 5 |
| 2/26/19 | 2018-2019 | Martinson | -29.88 | 2.79 | 30 |
| 2/28/19 | 2018-2019 | Londo | -24.86 | 2.21 | 5 |
| 3/4/19 | 2018-2019 | Martinson | -26.80 | 3.14 | 30 |
| 3/7/19 | 2018-2019 | Londo | -23.83 | 0.68 | 4 |
| 3/12/19 | 2018-2019 | Martinson | -28.86 | 3.50 | 30 |
| 3/13/19 | 2018-2019 | Londo | -21.90 | 1.87 | 10 |
| 3/18/19 | 2018-2019 | Martinson | -27.10 | 2.45 | 30 |
| 3/20/19 | 2018-2019 | Londo | -19.54 | 1.64 | 5 |
| 3/26/19 | 2018-2019 | Martinson | -25.53 | 3.06 | 30 |
| 3/28/19 | 2018-2019 | Londo | -17.48 | 1.22 | 5 |

| | | | | | |
|---|---|---|---|---|---|
| 4/2/19 | 2018-2019 | Martinson | -21.39 | 3.35 | 30 |
| 4/3/19 | 2018-2019 | Londo | -14.45 | 0.95 | 5 |
| 4/10/19 | 2018-2019 | Londo | -10.12 | 1.17 | 4 |
| 4/16/19 | 2018-2019 | Londo | -10.29 | 0.37 | 4 |
| 9/12/19 | 2019-2020 | Londo | -7.75 | 1.16 | 7 |
| 9/24/19 | 2019-2020 | Londo | -9.27 | 1.65 | 9 |
| 10/2/19 | 2019-2020 | Londo | -10.34 | 2.07 | 10 |
| 10/10/19 | 2019-2020 | Londo | -11.57 | 1.87 | 9 |
| 10/18/19 | 2019-2020 | Londo | -15.49 | 3.44 | 4 |
| 10/23/19 | 2019-2020 | Londo | -13.39 | 0.72 | 6 |
| 10/30/19 | 2019-2020 | Londo | -13.59 | 1.44 | 10 |
| 11/6/19 | 2019-2020 | Londo | -21.48 | 1.63 | 7 |
| 11/13/19 | 2019-2020 | Londo | -19.92 | 1.79 | 10 |
| 11/20/19 | 2019-2020 | Londo | -20.59 | 2.09 | 10 |
| 11/27/19 | 2019-2020 | Londo | -21.73 | 3.01 | 20 |
| 12/4/19 | 2019-2020 | Londo | -22.80 | 3.04 | 10 |
| 12/11/19 | 2019-2020 | Londo | -24.20 | 1.98 | 10 |
| 12/18/19 | 2019-2020 | Londo | -27.22 | 1.82 | 10 |
| 12/19/19 | 2019-2020 | Londo | -25.34 | 1.96 | 8 |
| 12/24/19 | 2019-2020 | Londo | -20.10 | 3.70 | 8 |
| 1/2/20 | 2019-2020 | Londo | -23.18 | 2.49 | 6 |
| 1/7/20 | 2019-2020 | Londo | -24.23 | 2.66 | 10 |
| 1/15/20 | 2019-2020 | Londo | -25.92 | 1.14 | 7 |
| 1/22/20 | 2019-2020 | Londo | -25.56 | 2.76 | 7 |
| 1/29/20 | 2019-2020 | Londo | -21.33 | 4.06 | 9 |
| 2/5/20 | 2019-2020 | Londo | -22.92 | 3.07 | 9 |
| 2/11/20 | 2019-2020 | Londo | -19.75 | 5.44 | 6 |
| 2/12/20 | 2019-2020 | Londo | -20.51 | 5.09 | 8 |
| 2/19/20 | 2019-2020 | Londo | -27.90 | 3.96 | 3 |
| 2/26/20 | 2019-2020 | Londo | -23.50 | 1.12 | 9 |
| 3/4/20 | 2019-2020 | Londo | -20.19 | 2.70 | 18 |
| 3/11/20 | 2019-2020 | Londo | -15.27 | 1.42 | 8 |
| 3/18/20 | 2019-2020 | Londo | -13.41 | 0.65 | 9 |
| 3/25/20 | 2019-2020 | Londo | -13.89 | 1.00 | 5 |
| 10/2/20 | 2020-2021 | Londo | -14.35 | 3.20 | 10 |
| 10/8/20 | 2020-2021 | Londo | -11.63 | 1.79 | 10 |
| 10/14/20 | 2020-2021 | Londo | -13.36 | 1.84 | 8 |
| 10/22/20 | 2020-2021 | Londo | -13.26 | 0.62 | 8 |
| 10/29/20 | 2020-2021 | Londo | -14.56 | 1.53 | 10 |
| 11/4/20 | 2020-2021 | Londo | -14.71 | 3.22 | 10 |

| | | | | | | |
|---|---|---|---|---|---|---|
| | 11/10/20 | 2020-2021 | Londo | -15.39 | 0.67 | 7 |
| | 11/18/20 | 2020-2021 | Londo | -20.30 | 2.12 | 18 |
| | 11/25/20 | 2020-2021 | Londo | -20.39 | 1.97 | 16 |
| | 12/2/20 | 2020-2021 | Londo | -20.67 | 0.75 | 7 |
| | 12/10/20 | 2020-2021 | Londo | -22.55 | 1.78 | 8 |
| | 12/15/20 | 2020-2021 | Londo | -21.58 | 2.46 | 7 |
| | 12/23/20 | 2020-2021 | Londo | -22.70 | 4.29 | 10 |
| | 12/30/20 | 2020-2021 | Londo | -26.37 | 1.94 | 10 |
| | 1/6/21 | 2020-2021 | Londo | -25.95 | 1.97 | 10 |
| | 1/13/21 | 2020-2021 | Londo | -25.54 | 2.90 | 10 |
| | 1/20/21 | 2020-2021 | Londo | -23.03 | 4.27 | 10 |
| | 1/27/21 | 2020-2021 | Londo | -25.83 | 1.19 | 8 |
| | 2/3/21 | 2020-2021 | Londo | -23.50 | 3.59 | 7 |
| | 2/10/21 | 2020-2021 | Londo | -25.66 | 1.23 | 9 |
| | 2/17/21 | 2020-2021 | Londo | -28.58 | 2.06 | 9 |
| | 2/24/21 | 2020-2021 | Londo | -27.41 | 1.72 | 10 |
| | 3/3/21 | 2020-2021 | Londo | -30.06 | 1.90 | 8 |
| | 3/10/21 | 2020-2021 | Londo | -26.36 | 4.23 | 9 |
| | 3/17/21 | 2020-2021 | Londo | -21.71 | 1.62 | 8 |
| | 3/24/21 | 2020-2021 | Londo | -18.10 | 0.90 | 10 |
| *V. vinifera* 'Cabernet Sauvignon' | 10/15/15 | 2015-2016 | Kovaleski | -9.74 | 1.92 | 8 |
| | 10/22/15 | 2015-2016 | Kovaleski | -11.30 | 1.51 | 7 |
| | 10/29/15 | 2015-2016 | Kovaleski | -14.63 | 2.18 | 7 |
| | 11/5/15 | 2015-2016 | Kovaleski | -11.37 | 1.70 | 7 |
| | 11/12/15 | 2015-2016 | Kovaleski | -16.59 | 1.01 | 8 |
| | 11/19/15 | 2015-2016 | Kovaleski | -13.08 | 4.16 | 5 |
| | 12/3/15 | 2015-2016 | Kovaleski | -18.75 | 1.23 | 4 |
| | 12/10/15 | 2015-2016 | Kovaleski | -21.18 | 1.09 | 5 |
| | 12/17/15 | 2015-2016 | Kovaleski | -21.59 | 0.65 | 7 |
| | 12/24/15 | 2015-2016 | Kovaleski | -22.00 | 1.75 | 8 |
| | 12/30/15 | 2015-2016 | Kovaleski | -18.82 | 4.02 | 5 |
| | 1/5/16 | 2015-2016 | Kovaleski | -23.10 | 0.90 | 5 |
| | 1/13/16 | 2015-2016 | Kovaleski | -21.67 | 2.43 | 6 |
| | 1/13/16 | 2015-2016 | Londo | -21.67 | 2.43 | 6 |
| | 1/18/16 | 2015-2016 | Londo | -25.20 | 0.34 | 3 |
| | 1/19/16 | 2015-2016 | Londo | -24.39 | 2.69 | 3 |
| | 1/20/16 | 2015-2016 | Londo | -21.84 | 2.49 | 2 |
| | 1/21/16 | 2015-2016 | Kovaleski | -21.74 | 3.10 | 5 |
| | 1/21/16 | 2015-2016 | Londo | -23.69 | 1.09 | 4 |
| | 1/22/16 | 2015-2016 | Londo | -23.79 | 1.41 | 4 |

| | | | | | |
|---|---|---|---|---|---|
| 1/23/16 | 2015-2016 | Londo | -24.64 | 0.43 | 3 |
| 1/24/16 | 2015-2016 | Londo | -22.25 | 1.03 | 4 |
| 1/25/16 | 2015-2016 | Londo | -23.76 | 2.30 | 5 |
| 1/26/16 | 2015-2016 | Londo | -22.03 | 2.57 | 5 |
| 1/27/16 | 2015-2016 | Londo | -20.97 | 3.91 | 4 |
| 1/28/16 | 2015-2016 | Kovaleski | -23.09 | 1.37 | 7 |
| 1/28/16 | 2015-2016 | Londo | -21.80 | 2.27 | 5 |
| 1/29/16 | 2015-2016 | Londo | -21.15 | 2.86 | 6 |
| 1/30/16 | 2015-2016 | Londo | -23.11 | 0.56 | 4 |
| 1/31/16 | 2015-2016 | Londo | -23.60 | 0.10 | 3 |
| 2/4/16 | 2015-2016 | Kovaleski | -24.65 | 2.80 | 8 |
| 2/11/16 | 2015-2016 | Kovaleski | -23.20 | 2.58 | 4 |
| 2/18/16 | 2015-2016 | Kovaleski | -22.48 | 2.25 | 4 |
| 2/26/16 | 2015-2016 | Kovaleski | -20.10 | 4.05 | 4 |
| 3/3/16 | 2015-2016 | Kovaleski | -22.66 | 0.93 | 7 |
| 3/10/16 | 2015-2016 | Kovaleski | -18.20 | 1.68 | 4 |
| 3/17/16 | 2015-2016 | Kovaleski | -17.28 | 3.09 | 4 |
| 3/24/16 | 2015-2016 | Kovaleski | -17.09 | 1.71 | 8 |
| 3/31/16 | 2015-2016 | Kovaleski | -18.25 | 0.64 | 2 |
| 4/7/16 | 2015-2016 | Kovaleski | -15.30 | 1.71 | 8 |
| 4/14/16 | 2015-2016 | Kovaleski | -15.10 | 1.35 | 4 |
| 4/21/16 | 2015-2016 | Kovaleski | -5.83 | 0.83 | 4 |
| 10/26/16 | 2016-2017 | Kovaleski | -10.32 | 2.61 | 5 |
| 11/1/16 | 2016-2017 | Kovaleski | -11.36 | 0.71 | 8 |
| 11/8/16 | 2016-2017 | Kovaleski | -12.60 | 1.06 | 8 |
| 11/15/16 | 2016-2017 | Kovaleski | -14.16 | 1.50 | 7 |
| 11/22/16 | 2016-2017 | Kovaleski | -17.54 | 0.55 | 8 |
| 11/29/16 | 2016-2017 | Kovaleski | -19.44 | 1.26 | 7 |
| 12/6/16 | 2016-2017 | Kovaleski | -20.00 | 0.62 | 8 |
| 12/13/16 | 2016-2017 | Kovaleski | -22.26 | 1.09 | 8 |
| 12/20/16 | 2016-2017 | Kovaleski | -23.91 | 0.54 | 8 |
| 12/27/16 | 2016-2017 | Kovaleski | -22.41 | 0.68 | 7 |
| 1/3/17 | 2016-2017 | Kovaleski | -23.40 | 0.99 | 6 |
| 1/10/17 | 2016-2017 | Kovaleski | -21.18 | 2.34 | 6 |
| 1/20/17 | 2016-2017 | Kovaleski | -22.37 | 0.85 | 7 |
| 1/25/17 | 2016-2017 | Kovaleski | -21.75 | 1.57 | 8 |
| 1/29/17 | 2016-2017 | Kovaleski | -19.77 | 2.26 | 6 |
| 2/8/17 | 2016-2017 | Kovaleski | -21.78 | 1.56 | 6 |
| 2/14/17 | 2016-2017 | Kovaleski | -23.41 | 1.33 | 7 |
| 2/20/17 | 2016-2017 | Kovaleski | -21.33 | 1.66 | 7 |

| | | | | | |
|---|---|---|---|---|---|
| 2/28/17 | 2016-2017 | Kovaleski | -20.11 | 1.19 | 7 |
| 3/7/17 | 2016-2017 | Kovaleski | -19.03 | 2.67 | 6 |
| 3/14/17 | 2016-2017 | Kovaleski | -22.09 | 1.07 | 8 |
| 3/21/17 | 2016-2017 | Kovaleski | -19.66 | 2.62 | 5 |
| 3/28/17 | 2016-2017 | Kovaleski | -19.83 | 0.57 | 7 |
| 4/4/17 | 2016-2017 | Kovaleski | -16.19 | 3.38 | 7 |
| 4/11/17 | 2016-2017 | Kovaleski | -11.13 | 3.68 | 6 |
| 4/18/17 | 2016-2017 | Kovaleski | -6.16 | 3.00 | 7 |
| 10/6/17 | 2017-2018 | Kovaleski | -9.68 | 0.02 | 2 |
| 10/13/17 | 2017-2018 | Kovaleski | -8.58 | 1.62 | 5 |
| 10/20/17 | 2017-2018 | Kovaleski | -10.17 | 1.61 | 6 |
| 10/27/17 | 2017-2018 | Kovaleski | -9.72 | 1.52 | 5 |
| 11/3/17 | 2017-2018 | Kovaleski | -11.22 | 2.27 | 5 |
| 11/11/17 | 2017-2018 | Kovaleski | -13.36 | 1.80 | 5 |
| 11/17/17 | 2017-2018 | Kovaleski | -17.48 | 0.52 | 5 |
| 11/24/17 | 2017-2018 | Kovaleski | -17.50 | 0.72 | 6 |
| 12/1/17 | 2017-2018 | Kovaleski | -17.48 | 0.44 | 5 |
| 12/8/17 | 2017-2018 | Kovaleski | -18.32 | 0.63 | 5 |
| 12/19/17 | 2017-2018 | Kovaleski | -19.78 | 0.43 | 4 |
| 12/26/17 | 2017-2018 | Kovaleski | -20.66 | 1.15 | 5 |
| 1/2/18 | 2017-2018 | Kovaleski | -22.55 | 0.53 | 4 |
| 1/12/18 | 2017-2018 | Kovaleski | -20.78 | 0.57 | 4 |
| 1/19/18 | 2017-2018 | Kovaleski | -20.36 | 1.14 | 5 |
| 1/26/18 | 2017-2018 | Kovaleski | -23.90 | 0.95 | 5 |
| 1/31/18 | 2017-2018 | Kovaleski | -22.43 | 0.77 | 6 |
| 2/8/18 | 2017-2018 | Kovaleski | -20.80 | 1.24 | 4 |
| 2/22/18 | 2017-2018 | Kovaleski | -21.50 | 1.77 | 5 |
| 3/1/18 | 2017-2018 | Kovaleski | -21.77 | 1.07 | 3 |
| 3/8/18 | 2017-2018 | Kovaleski | -21.56 | 1.43 | 5 |
| 3/15/18 | 2017-2018 | Kovaleski | -20.90 | 1.54 | 4 |
| 3/22/18 | 2017-2018 | Kovaleski | -21.20 | 0.57 | 2 |
| 3/30/18 | 2017-2018 | Kovaleski | -22.75 | 0.50 | 4 |
| 4/2/18 | 2017-2018 | Kovaleski | -20.50 | 1.56 | 6 |
| 4/5/18 | 2017-2018 | Kovaleski | -18.23 | 1.30 | 3 |
| 4/12/18 | 2017-2018 | Kovaleski | -19.23 | 2.24 | 4 |
| 4/20/18 | 2017-2018 | Kovaleski | -18.57 | 1.04 | 3 |
| 4/26/18 | 2017-2018 | Kovaleski | -13.58 | 3.80 | 4 |
| 5/3/18 | 2017-2018 | Kovaleski | -7.08 | 2.32 | 6 |
| 9/20/18 | 2018-2019 | Londo | -5.03 | 0.95 | 3 |
| 9/27/18 | 2018-2019 | Londo | -5.63 | 0.61 | 2 |

| | | | | | |
|---|---|---|---|---|---|
| 10/3/18 | 2018-2019 | Londo | -6.30 | 1.23 | 4 |
| 10/9/18 | 2018-2019 | Londo | -8.78 | 0.69 | 5 |
| 10/17/18 | 2018-2019 | Londo | -8.53 | 2.79 | 4 |
| 10/24/18 | 2018-2019 | Londo | -8.88 | 2.06 | 4 |
| 10/30/18 | 2018-2019 | Londo | -10.60 | 0.28 | 2 |
| 11/7/18 | 2018-2019 | Londo | -11.84 | 1.18 | 5 |
| 11/8/18 | 2018-2019 | Londo | -11.15 | 1.52 | 4 |
| 11/15/18 | 2018-2019 | Londo | -14.53 | 2.92 | 4 |
| 11/16/18 | 2018-2019 | Londo | -15.78 | 2.05 | 4 |
| 11/23/18 | 2018-2019 | Londo | -16.35 | 0.71 | 4 |
| 11/29/18 | 2018-2019 | Londo | -17.27 | 0.95 | 3 |
| 12/6/18 | 2018-2019 | Londo | -17.83 | 2.57 | 3 |
| 12/13/18 | 2018-2019 | Londo | -18.43 | 4.51 | 4 |
| 12/19/18 | 2018-2019 | Londo | -19.83 | 0.53 | 4 |
| 12/27/18 | 2018-2019 | Londo | -19.49 | 1.44 | 5 |
| 1/4/19 | 2018-2019 | Londo | -19.12 | 2.53 | 5 |
| 1/11/19 | 2018-2019 | Londo | -18.42 | 2.51 | 5 |
| 1/16/19 | 2018-2019 | Londo | -22.05 | 1.55 | 4 |
| 1/25/19 | 2018-2019 | Londo | -20.89 | 1.95 | 4 |
| 1/29/19 | 2018-2019 | Londo | -19.62 | 1.99 | 5 |
| 2/4/19 | 2018-2019 | Londo | -20.56 | 2.01 | 5 |
| 2/11/19 | 2018-2019 | Londo | -18.35 | 3.93 | 8 |
| 2/21/19 | 2018-2019 | Londo | -19.32 | 2.13 | 5 |
| 2/28/19 | 2018-2019 | Londo | -20.93 | 1.40 | 3 |
| 3/7/19 | 2018-2019 | Londo | -20.93 | 1.60 | 3 |
| 3/13/19 | 2018-2019 | Londo | -21.09 | 1.31 | 4 |
| 3/20/19 | 2018-2019 | Londo | -20.17 | 1.73 | 4 |
| 3/28/19 | 2018-2019 | Londo | -18.50 | 0.95 | 3 |
| 4/3/19 | 2018-2019 | Londo | -17.69 | 0.87 | 4 |
| 4/10/19 | 2018-2019 | Londo | -14.53 | 1.11 | 3 |
| 4/16/19 | 2018-2019 | Londo | -11.06 | 2.12 | 5 |
| 9/12/19 | 2019-2020 | Londo | -6.47 | 1.14 | 3 |
| 9/24/19 | 2019-2020 | Londo | -8.77 | 1.46 | 4 |
| 10/2/19 | 2019-2020 | Londo | -8.09 | 1.04 | 4 |
| 10/10/19 | 2019-2020 | Londo | -7.84 | 1.12 | 7 |
| 10/18/19 | 2019-2020 | Londo | -8.95 | 0.79 | 6 |
| 10/23/19 | 2019-2020 | Londo | -10.33 | 0.76 | 8 |
| 10/30/19 | 2019-2020 | Londo | -10.06 | 1.18 | 8 |
| 11/6/19 | 2019-2020 | Londo | -13.24 | 2.61 | 5 |
| 11/13/19 | 2019-2020 | Londo | -13.81 | 2.13 | 9 |

| | | | | | |
|---|---|---|---|---|---|
| 11/20/19 | 2019-2020 | Londo | -17.34 | 0.95 | 10 |
| 11/27/19 | 2019-2020 | Londo | -18.22 | 1.36 | 20 |
| 12/4/19 | 2019-2020 | Londo | -19.70 | 0.82 | 9 |
| 12/11/19 | 2019-2020 | Londo | -19.84 | 3.28 | 4 |
| 12/18/19 | 2019-2020 | Londo | -19.90 | 1.37 | 7 |
| 12/19/19 | 2019-2020 | Londo | -21.69 | 1.77 | 5 |
| 12/24/19 | 2019-2020 | Londo | -18.93 | 2.37 | 6 |
| 1/2/20 | 2019-2020 | Londo | -19.58 | 2.94 | 8 |
| 1/7/20 | 2019-2020 | Londo | -22.51 | 0.73 | 10 |
| 1/15/20 | 2019-2020 | Londo | -20.66 | 1.03 | 8 |
| 1/22/20 | 2019-2020 | Londo | -19.93 | 2.35 | 7 |
| 1/29/20 | 2019-2020 | Londo | -18.81 | 2.75 | 4 |
| 2/5/20 | 2019-2020 | Londo | -19.24 | 1.81 | 4 |
| 2/11/20 | 2019-2020 | Londo | -15.86 | 3.06 | 6 |
| 2/12/20 | 2019-2020 | Londo | -20.11 | 3.96 | 7 |
| 2/19/20 | 2019-2020 | Londo | -24.55 | 2.32 | 3 |
| 2/26/20 | 2019-2020 | Londo | -21.20 | 1.82 | 10 |
| 3/4/20 | 2019-2020 | Londo | -18.62 | 1.92 | 8 |
| 3/11/20 | 2019-2020 | Londo | -18.19 | 1.64 | 2 |
| 3/18/20 | 2019-2020 | Londo | -17.37 | 1.40 | 7 |
| 3/25/20 | 2019-2020 | Londo | -15.73 | 0.46 | 4 |
| 10/2/20 | 2020-2021 | Londo | -12.35 | 0.44 | 6 |
| 10/8/20 | 2020-2021 | Londo | -8.55 | 0.46 | 5 |
| 10/14/20 | 2020-2021 | Londo | -10.30 | 1.30 | 7 |
| 10/22/20 | 2020-2021 | Londo | -9.65 | 1.54 | 6 |
| 10/29/20 | 2020-2021 | Londo | -10.38 | 1.58 | 10 |
| 11/4/20 | 2020-2021 | Londo | -14.10 | 3.29 | 9 |
| 11/10/20 | 2020-2021 | Londo | -12.38 | 1.34 | 10 |
| 11/18/20 | 2020-2021 | Londo | -15.87 | 1.12 | 18 |
| 11/25/20 | 2020-2021 | Londo | -17.09 | 1.61 | 12 |
| 12/2/20 | 2020-2021 | Londo | -17.65 | 2.28 | 9 |
| 12/10/20 | 2020-2021 | Londo | -19.97 | 1.79 | 6 |
| 12/15/20 | 2020-2021 | Londo | -21.55 | 1.40 | 6 |
| 12/23/20 | 2020-2021 | Londo | -20.86 | 1.43 | 8 |
| 12/30/20 | 2020-2021 | Londo | -22.16 | 2.79 | 6 |
| 1/6/21 | 2020-2021 | Londo | -22.41 | 0.96 | 5 |
| 1/13/21 | 2020-2021 | Londo | -21.85 | 2.71 | 6 |
| 1/20/21 | 2020-2021 | Londo | -21.27 | 2.98 | 7 |
| 1/27/21 | 2020-2021 | Londo | -22.21 | 1.70 | 8 |
| 2/3/21 | 2020-2021 | Londo | -22.70 | 1.74 | 6 |

| | | | | | | |
|---|---|---|---|---|---|---|
| | 2/10/21 | 2020-2021 | Londo | -20.60 | 2.69 | 9 |
| | 2/17/21 | 2020-2021 | Londo | -21.07 | 2.99 | 9 |
| | 2/24/21 | 2020-2021 | Londo | -21.67 | 3.60 | 4 |
| | 3/3/21 | 2020-2021 | Londo | -20.57 | 3.21 | 7 |
| | 3/10/21 | 2020-2021 | Londo | -19.79 | 2.19 | 5 |
| | 3/17/21 | 2020-2021 | Londo | -19.19 | 2.36 | 10 |
| | 3/24/21 | 2020-2021 | Londo | -15.56 | 1.09 | 5 |
| *V. vinifera* ‘Riesling’ | 10/31/12 | 2012-2013 | Martinson | -13.92 | 0.76 | 30 |
| | 11/6/12 | 2012-2013 | Martinson | -15.62 | 1.14 | 30 |
| | 11/13/12 | 2012-2013 | Martinson | -19.89 | 1.01 | 30 |
| | 11/20/12 | 2012-2013 | Martinson | -21.93 | 0.74 | 30 |
| | 11/27/12 | 2012-2013 | Martinson | -21.84 | 0.96 | 30 |
| | 12/4/12 | 2012-2013 | Martinson | -22.61 | 1.10 | 30 |
| | 12/11/12 | 2012-2013 | Martinson | -22.10 | 1.29 | 30 |
| | 12/18/12 | 2012-2013 | Martinson | -22.52 | 1.40 | 30 |
| | 12/26/12 | 2012-2013 | Martinson | -24.50 | 1.19 | 30 |
| | 1/2/13 | 2012-2013 | Martinson | -25.57 | 1.72 | 30 |
| | 1/9/13 | 2012-2013 | Martinson | -25.38 | 1.29 | 30 |
| | 1/15/13 | 2012-2013 | Martinson | -23.53 | 1.26 | 30 |
| | 1/22/13 | 2012-2013 | Martinson | -25.51 | 1.69 | 30 |
| | 1/28/13 | 2012-2013 | Martinson | -24.83 | 1.42 | 30 |
| | 2/4/13 | 2012-2013 | Martinson | -25.14 | 2.15 | 30 |
| | 2/11/13 | 2012-2013 | Martinson | -24.31 | 1.76 | 30 |
| | 2/18/13 | 2012-2013 | Martinson | -25.93 | 1.42 | 30 |
| | 2/26/13 | 2012-2013 | Martinson | -23.38 | 1.75 | 30 |
| | 3/4/13 | 2012-2013 | Martinson | -24.26 | 1.08 | 30 |
| | 3/11/13 | 2012-2013 | Martinson | -22.95 | 0.98 | 30 |
| | 3/18/13 | 2012-2013 | Martinson | -22.73 | 1.20 | 30 |
| | 11/4/13 | 2013-2014 | Martinson | -17.55 | 1.52 | 27 |
| | 11/11/13 | 2013-2014 | Martinson | -19.67 | 0.83 | 27 |
| | 11/25/13 | 2013-2014 | Martinson | -22.95 | 1.06 | 27 |
| | 12/2/13 | 2013-2014 | Martinson | -22.66 | 0.74 | 27 |
| | 12/9/13 | 2013-2014 | Martinson | -22.90 | 1.15 | 27 |
| | 12/16/13 | 2013-2014 | Martinson | -24.12 | 1.39 | 27 |
| | 12/23/13 | 2013-2014 | Martinson | -23.07 | 1.10 | 27 |
| | 12/30/13 | 2013-2014 | Martinson | -23.74 | 0.87 | 27 |
| | 1/6/14 | 2013-2014 | Martinson | -24.53 | 1.31 | 27 |
| | 1/13/14 | 2013-2014 | Martinson | -23.72 | 1.70 | 27 |
| | 1/20/14 | 2013-2014 | Martinson | -24.57 | 0.96 | 27 |
| | 2/4/14 | 2013-2014 | Martinson | -25.35 | 1.09 | 27 |

| | | | | | |
|---|---|---|---|---|---|
| 2/10/14 | 2013-2014 | Martinson | -25.21 | 1.61 | 27 |
| 2/17/14 | 2013-2014 | Martinson | -25.40 | 1.29 | 27 |
| 2/24/14 | 2013-2014 | Martinson | -24.95 | 1.02 | 27 |
| 3/3/14 | 2013-2014 | Martinson | -25.18 | 1.99 | 23 |
| 3/10/14 | 2013-2014 | Martinson | -24.90 | 1.61 | 27 |
| 3/18/14 | 2013-2014 | Martinson | -24.46 | 0.96 | 45 |
| 3/24/14 | 2013-2014 | Martinson | -24.10 | 0.88 | 36 |
| 3/31/14 | 2013-2014 | Martinson | -22.50 | 0.84 | 41 |
| 4/7/14 | 2013-2014 | Martinson | -21.01 | 0.81 | 49 |
| 4/15/14 | 2013-2014 | Martinson | -13.39 | 2.20 | 45 |
| 11/4/14 | 2014-2015 | Martinson | -14.42 | 1.02 | 60 |
| 11/12/14 | 2014-2015 | Martinson | -17.89 | 1.23 | 60 |
| 11/17/14 | 2014-2015 | Martinson | -20.96 | 1.03 | 60 |
| 11/25/14 | 2014-2015 | Martinson | -21.04 | 0.95 | 60 |
| 12/1/14 | 2014-2015 | Martinson | -22.15 | 0.88 | 60 |
| 12/8/14 | 2014-2015 | Martinson | -21.86 | 1.78 | 60 |
| 12/15/14 | 2014-2015 | Martinson | -22.91 | 1.73 | 60 |
| 12/18/14 | 2014-2015 | Kovaleski | -22.95 | 1.91 | 7 |
| 12/22/14 | 2014-2015 | Martinson | -23.54 | 0.78 | 60 |
| 12/29/14 | 2014-2015 | Martinson | -23.01 | 1.05 | 60 |
| 1/5/15 | 2014-2015 | Martinson | -23.76 | 3.79 | 60 |
| 1/12/15 | 2014-2015 | Martinson | -24.68 | 1.09 | 60 |
| 1/19/15 | 2014-2015 | Martinson | -24.51 | 1.50 | 60 |
| 1/26/15 | 2014-2015 | Martinson | -24.71 | 2.12 | 60 |
| 2/3/15 | 2014-2015 | Martinson | -26.16 | 2.02 | 30 |
| 2/10/15 | 2014-2015 | Martinson | -24.87 | 2.01 | 60 |
| 2/11/15 | 2014-2015 | Kovaleski | -25.49 | 1.44 | 8 |
| 2/16/15 | 2014-2015 | Martinson | -26.65 | 3.15 | 29 |
| 2/17/15 | 2014-2015 | Martinson | -25.83 | 5.46 | 30 |
| 2/24/15 | 2014-2015 | Martinson | -25.45 | 3.08 | 50 |
| 3/2/15 | 2014-2015 | Martinson | -26.71 | 2.20 | 60 |
| 3/9/15 | 2014-2015 | Martinson | -25.13 | 1.91 | 60 |
| 3/16/15 | 2014-2015 | Martinson | -25.15 | 1.99 | 60 |
| 3/23/15 | 2014-2015 | Martinson | -24.66 | 2.29 | 60 |
| 3/30/15 | 2014-2015 | Martinson | -24.32 | 2.02 | 60 |
| 4/6/15 | 2014-2015 | Martinson | -22.05 | 1.37 | 60 |
| 10/15/15 | 2015-2016 | Kovaleski | -12.11 | 1.71 | 16 |
| 10/22/15 | 2015-2016 | Kovaleski | -12.35 | 1.64 | 14 |
| 10/29/15 | 2015-2016 | Kovaleski | -17.40 | 3.71 | 14 |
| 11/5/15 | 2015-2016 | Kovaleski | -14.42 | 1.31 | 15 |

| | | | | | |
|---|---|---|---|---|---|
| 11/12/15 | 2015-2016 | Kovaleski | -18.69 | 2.55 | 14 |
| 11/19/15 | 2015-2016 | Kovaleski | -18.46 | 1.65 | 10 |
| 11/24/15 | 2015-2016 | Kovaleski | -21.21 | 4.39 | 15 |
| 12/3/15 | 2015-2016 | Kovaleski | -21.71 | 1.45 | 13 |
| 12/10/15 | 2015-2016 | Kovaleski | -22.69 | 1.33 | 13 |
| 12/17/15 | 2015-2016 | Kovaleski | -22.30 | 1.65 | 12 |
| 12/24/15 | 2015-2016 | Kovaleski | -22.59 | 1.67 | 13 |
| 12/30/15 | 2015-2016 | Kovaleski | -18.30 | 6.70 | 10 |
| 1/5/16 | 2015-2016 | Kovaleski | -22.68 | 6.63 | 6 |
| 1/13/16 | 2015-2016 | Kovaleski | -25.88 | 1.64 | 11 |
| 1/13/16 | 2015-2016 | Londo | -25.30 | 1.28 | 6 |
| 1/18/16 | 2015-2016 | Londo | -25.45 | 0.93 | 3 |
| 1/19/16 | 2015-2016 | Londo | -26.09 | 1.04 | 4 |
| 1/20/16 | 2015-2016 | Londo | -24.87 | 1.80 | 3 |
| 1/21/16 | 2015-2016 | Kovaleski | -26.32 | 1.10 | 10 |
| 1/21/16 | 2015-2016 | Londo | -24.29 | 0.42 | 3 |
| 1/22/16 | 2015-2016 | Londo | -24.89 | 1.95 | 3 |
| 1/23/16 | 2015-2016 | Londo | -24.80 | 2.26 | 4 |
| 1/24/16 | 2015-2016 | Londo | -25.40 | 2.43 | 4 |
| 1/25/16 | 2015-2016 | Londo | -26.00 | 1.14 | 5 |
| 1/26/16 | 2015-2016 | Londo | -25.15 | 2.15 | 6 |
| 1/27/16 | 2015-2016 | Londo | -23.57 | 1.62 | 3 |
| 1/28/16 | 2015-2016 | Kovaleski | -24.20 | 1.99 | 15 |
| 1/28/16 | 2015-2016 | Londo | -25.25 | 0.19 | 3 |
| 1/29/16 | 2015-2016 | Londo | -23.46 | 1.82 | 4 |
| 1/30/16 | 2015-2016 | Londo | -25.27 | 1.18 | 4 |
| 1/31/16 | 2015-2016 | Londo | -22.64 | 2.40 | 5 |
| 2/4/16 | 2015-2016 | Kovaleski | -24.57 | 1.34 | 15 |
| 2/11/16 | 2015-2016 | Kovaleski | -25.54 | 1.67 | 10 |
| 2/18/16 | 2015-2016 | Kovaleski | -25.30 | 2.07 | 12 |
| 2/26/16 | 2015-2016 | Kovaleski | -24.68 | 1.11 | 12 |
| 3/3/16 | 2015-2016 | Kovaleski | -25.29 | 0.94 | 7 |
| 3/10/16 | 2015-2016 | Kovaleski | -20.76 | 1.02 | 9 |
| 3/17/16 | 2015-2016 | Kovaleski | -17.86 | 1.60 | 13 |
| 3/24/16 | 2015-2016 | Kovaleski | -19.33 | 1.71 | 7 |
| 3/31/16 | 2015-2016 | Kovaleski | -13.46 | 1.82 | 10 |
| 4/7/16 | 2015-2016 | Kovaleski | -13.46 | 1.72 | 12 |
| 4/14/16 | 2015-2016 | Kovaleski | -12.83 | 2.80 | 15 |
| 4/21/16 | 2015-2016 | Kovaleski | -5.10 | 2.10 | 4 |
| 10/4/16 | 2016-2017 | Kovaleski | -11.78 | 0.98 | 8 |

| | | | | | |
|---|---|---|---|---|---|
| 10/11/16 | 2016-2017 | Kovaleski | -13.30 | 2.50 | 8 |
| 10/20/16 | 2016-2017 | Kovaleski | -12.97 | 0.26 | 7 |
| 10/26/16 | 2016-2017 | Kovaleski | -12.13 | 1.82 | 7 |
| 11/1/16 | 2016-2017 | Kovaleski | -13.80 | 1.06 | 7 |
| 11/1/16 | 2016-2017 | Martinson | -12.51 | 1.35 | 30 |
| 11/8/16 | 2016-2017 | Kovaleski | -15.32 | 0.61 | 6 |
| 11/8/16 | 2016-2017 | Martinson | -12.47 | 1.82 | 30 |
| 11/14/16 | 2016-2017 | Martinson | -16.79 | 0.78 | 30 |
| 11/15/16 | 2016-2017 | Kovaleski | -17.44 | 1.11 | 8 |
| 11/22/16 | 2016-2017 | Kovaleski | -17.85 | 1.10 | 8 |
| 11/22/16 | 2016-2017 | Martinson | -19.20 | 1.39 | 30 |
| 11/28/16 | 2016-2017 | Martinson | -21.02 | 1.69 | 30 |
| 11/29/16 | 2016-2017 | Kovaleski | -20.29 | 0.97 | 8 |
| 12/5/16 | 2016-2017 | Martinson | -20.57 | 1.76 | 30 |
| 12/6/16 | 2016-2017 | Kovaleski | -19.98 | 0.63 | 8 |
| 12/12/16 | 2016-2017 | Martinson | -23.60 | 1.81 | 30 |
| 12/13/16 | 2016-2017 | Kovaleski | -23.32 | 1.17 | 6 |
| 12/19/16 | 2016-2017 | Martinson | -25.41 | 1.52 | 30 |
| 12/20/16 | 2016-2017 | Kovaleski | -24.70 | 0.48 | 8 |
| 12/26/16 | 2016-2017 | Martinson | -24.72 | 1.59 | 30 |
| 12/27/16 | 2016-2017 | Kovaleski | -23.21 | 1.00 | 8 |
| 1/2/17 | 2016-2017 | Martinson | -26.21 | 1.45 | 30 |
| 1/3/17 | 2016-2017 | Kovaleski | -24.86 | 1.05 | 7 |
| 1/10/17 | 2016-2017 | Kovaleski | -25.34 | 0.79 | 7 |
| 1/11/17 | 2016-2017 | Martinson | -24.56 | 1.24 | 30 |
| 1/16/17 | 2016-2017 | Martinson | -25.59 | 1.72 | 30 |
| 1/20/17 | 2016-2017 | Kovaleski | -23.80 | 1.09 | 7 |
| 1/23/17 | 2016-2017 | Martinson | -23.16 | 1.39 | 30 |
| 1/25/17 | 2016-2017 | Kovaleski | -23.58 | 0.81 | 6 |
| 1/29/17 | 2016-2017 | Kovaleski | -23.46 | 1.01 | 8 |
| 1/30/17 | 2016-2017 | Martinson | -25.75 | 1.38 | 30 |
| 2/6/17 | 2016-2017 | Martinson | -25.98 | 1.16 | 30 |
| 2/8/17 | 2016-2017 | Kovaleski | -22.21 | 1.72 | 8 |
| 2/14/17 | 2016-2017 | Kovaleski | -23.75 | 1.30 | 8 |
| 2/14/17 | 2016-2017 | Martinson | -24.88 | 1.84 | 30 |
| 2/20/17 | 2016-2017 | Kovaleski | -23.45 | 0.52 | 6 |
| 2/20/17 | 2016-2017 | Martinson | -22.95 | 1.10 | 30 |
| 2/27/17 | 2016-2017 | Martinson | -21.91 | 2.44 | 30 |
| 2/28/17 | 2016-2017 | Kovaleski | -19.44 | 1.14 | 5 |
| 3/7/17 | 2016-2017 | Kovaleski | -20.11 | 1.12 | 7 |

| | | | | | |
|---|---|---|---|---|---|
| 3/13/17 | 2016-2017 | Martinson | -23.70 | 1.74 | 30 |
| 3/14/17 | 2016-2017 | Kovaleski | -21.29 | 0.90 | 7 |
| 3/20/17 | 2016-2017 | Martinson | -20.07 | 1.64 | 30 |
| 3/21/17 | 2016-2017 | Kovaleski | -20.27 | 1.47 | 6 |
| 3/27/17 | 2016-2017 | Martinson | -20.97 | 1.29 | 30 |
| 3/28/17 | 2016-2017 | Kovaleski | -18.49 | 0.70 | 8 |
| 4/3/17 | 2016-2017 | Martinson | -19.51 | 1.44 | 30 |
| 4/4/17 | 2016-2017 | Kovaleski | -17.06 | 0.80 | 7 |
| 4/10/17 | 2016-2017 | Martinson | -13.39 | 1.65 | 30 |
| 4/11/17 | 2016-2017 | Kovaleski | -8.86 | 2.37 | 8 |
| 4/18/17 | 2016-2017 | Kovaleski | -4.86 | 1.09 | 7 |
| 10/6/17 | 2017-2018 | Kovaleski | -10.70 | 1.90 | 5 |
| 10/13/17 | 2017-2018 | Kovaleski | -12.50 | 0.99 | 4 |
| 10/20/17 | 2017-2018 | Kovaleski | -12.46 | 0.43 | 5 |
| 10/27/17 | 2017-2018 | Kovaleski | -13.10 | 0.55 | 4 |
| 11/3/17 | 2017-2018 | Kovaleski | -14.95 | 2.94 | 6 |
| 11/7/17 | 2017-2018 | Martinson | -14.48 | 1.64 | 30 |
| 11/11/17 | 2017-2018 | Kovaleski | -16.75 | 0.85 | 6 |
| 11/13/17 | 2017-2018 | Martinson | -17.47 | 2.02 | 30 |
| 11/17/17 | 2017-2018 | Kovaleski | -19.43 | 0.31 | 6 |
| 11/20/17 | 2017-2018 | Martinson | -19.71 | 1.67 | 30 |
| 11/24/17 | 2017-2018 | Kovaleski | -19.08 | 1.40 | 5 |
| 11/28/17 | 2017-2018 | Martinson | -20.81 | 0.85 | 30 |
| 12/1/17 | 2017-2018 | Kovaleski | -20.52 | 1.34 | 6 |
| 12/4/17 | 2017-2018 | Martinson | -21.24 | 1.55 | 30 |
| 12/8/17 | 2017-2018 | Kovaleski | -20.40 | 1.19 | 6 |
| 12/12/17 | 2017-2018 | Martinson | -22.45 | 1.56 | 30 |
| 12/18/17 | 2017-2018 | Martinson | -23.75 | 1.51 | 30 |
| 12/19/17 | 2017-2018 | Kovaleski | -22.06 | 1.01 | 5 |
| 12/26/17 | 2017-2018 | Kovaleski | -23.85 | 0.55 | 4 |
| 1/2/18 | 2017-2018 | Kovaleski | -21.64 | 1.33 | 5 |
| 1/3/18 | 2017-2018 | Martinson | -26.22 | 3.69 | 30 |
| 1/8/18 | 2017-2018 | Martinson | -23.51 | 2.87 | 30 |
| 1/12/18 | 2017-2018 | Kovaleski | -20.86 | 1.76 | 5 |
| 1/16/18 | 2017-2018 | Martinson | -23.98 | 2.31 | 30 |
| 1/19/18 | 2017-2018 | Kovaleski | -22.72 | 1.03 | 5 |
| 1/24/18 | 2017-2018 | Martinson | -22.74 | 1.33 | 30 |
| 1/26/18 | 2017-2018 | Kovaleski | -25.57 | 0.80 | 6 |
| 1/29/18 | 2017-2018 | Martinson | -22.30 | 1.51 | 30 |
| 1/31/18 | 2017-2018 | Kovaleski | -23.25 | 1.38 | 4 |

| | | | | | |
|---|---|---|---|---|---|
| 2/5/18 | 2017-2018 | Martinson | -22.84 | 1.85 | 30 |
| 2/8/18 | 2017-2018 | Kovaleski | -23.05 | 1.13 | 6 |
| 2/14/18 | 2017-2018 | Martinson | -23.49 | 1.40 | 30 |
| 2/20/18 | 2017-2018 | Martinson | -22.71 | 2.31 | 30 |
| 2/22/18 | 2017-2018 | Kovaleski | -22.65 | 0.93 | 4 |
| 2/26/18 | 2017-2018 | Martinson | -21.09 | 1.14 | 30 |
| 3/1/18 | 2017-2018 | Kovaleski | -23.38 | 1.53 | 5 |
| 3/5/18 | 2017-2018 | Martinson | -21.38 | 1.98 | 30 |
| 3/8/18 | 2017-2018 | Kovaleski | -22.62 | 1.12 | 6 |
| 3/12/18 | 2017-2018 | Martinson | -21.27 | 1.80 | 30 |
| 3/15/18 | 2017-2018 | Kovaleski | -22.40 | 0.44 | 3 |
| 3/19/18 | 2017-2018 | Martinson | -23.04 | 2.23 | 30 |
| 3/22/18 | 2017-2018 | Kovaleski | -22.14 | 1.95 | 5 |
| 3/26/18 | 2017-2018 | Martinson | -22.51 | 2.92 | 30 |
| 3/30/18 | 2017-2018 | Kovaleski | -19.95 | 0.35 | 2 |
| 4/2/18 | 2017-2018 | Kovaleski | -19.22 | 2.04 | 6 |
| 4/2/18 | 2017-2018 | Martinson | -19.70 | 1.69 | 30 |
| 4/5/18 | 2017-2018 | Kovaleski | -19.37 | 0.67 | 3 |
| 4/9/18 | 2017-2018 | Martinson | -21.36 | 1.45 | 30 |
| 4/12/18 | 2017-2018 | Kovaleski | -18.78 | 2.59 | 6 |
| 4/20/18 | 2017-2018 | Kovaleski | -17.68 | 1.09 | 4 |
| 4/26/18 | 2017-2018 | Kovaleski | -6.73 | 1.25 | 3 |
| 5/3/18 | 2017-2018 | Kovaleski | -4.55 | 1.06 | 2 |
| 9/20/18 | 2018-2019 | Londo | -8.76 | 0.79 | 5 |
| 9/27/18 | 2018-2019 | Londo | -9.28 | 1.22 | 4 |
| 10/3/18 | 2018-2019 | Londo | -10.80 | 0.55 | 4 |
| 10/9/18 | 2018-2019 | Londo | -11.42 | 0.58 | 5 |
| 10/16/18 | 2018-2019 | Martinson | -12.17 | 1.49 | 30 |
| 10/17/18 | 2018-2019 | Londo | -11.84 | 0.78 | 5 |
| 10/22/18 | 2018-2019 | Martinson | -13.50 | 2.70 | 30 |
| 10/24/18 | 2018-2019 | Londo | -12.42 | 2.20 | 5 |
| 10/29/18 | 2018-2019 | Martinson | -13.66 | 1.42 | 30 |
| 10/30/18 | 2018-2019 | Londo | -11.53 | 1.55 | 3 |
| 11/5/18 | 2018-2019 | Martinson | -15.84 | 2.06 | 30 |
| 11/7/18 | 2018-2019 | Londo | -15.40 | 0.96 | 3 |
| 11/8/18 | 2018-2019 | Londo | -15.34 | 0.77 | 5 |
| 11/12/18 | 2018-2019 | Martinson | -18.40 | 1.76 | 30 |
| 11/15/18 | 2018-2019 | Londo | -17.23 | 0.87 | 4 |
| 11/16/18 | 2018-2019 | Londo | -18.35 | 0.92 | 4 |
| 11/19/18 | 2018-2019 | Martinson | -21.35 | 1.68 | 30 |

| | | | | | |
|---|---|---|---|---|---|
| 11/23/18 | 2018-2019 | Londo | -17.88 | 0.95 | 5 |
| 11/26/18 | 2018-2019 | Martinson | -20.59 | 1.97 | 30 |
| 11/29/18 | 2018-2019 | Londo | -19.47 | 0.31 | 3 |
| 12/3/18 | 2018-2019 | Martinson | -21.51 | 1.52 | 30 |
| 12/6/18 | 2018-2019 | Londo | -21.32 | 1.36 | 5 |
| 12/12/18 | 2018-2019 | Martinson | -22.79 | 1.65 | 30 |
| 12/13/18 | 2018-2019 | Londo | -22.93 | 0.32 | 3 |
| 12/17/18 | 2018-2019 | Martinson | -22.00 | 1.44 | 30 |
| 12/19/18 | 2018-2019 | Londo | -22.28 | 0.42 | 10 |
| 12/27/18 | 2018-2019 | Londo | -21.32 | 0.63 | 5 |
| 1/3/19 | 2018-2019 | Martinson | -21.38 | 1.98 | 30 |
| 1/4/19 | 2018-2019 | Londo | -22.85 | 0.84 | 5 |
| 1/8/19 | 2018-2019 | Martinson | -21.38 | 1.98 | 30 |
| 1/11/19 | 2018-2019 | Londo | -23.57 | 1.28 | 5 |
| 1/14/19 | 2018-2019 | Martinson | -24.75 | 2.15 | 30 |
| 1/16/19 | 2018-2019 | Londo | -23.34 | 0.64 | 5 |
| 1/21/19 | 2018-2019 | Martinson | -25.61 | 1.70 | 30 |
| 1/25/19 | 2018-2019 | Londo | -23.00 | 1.20 | 5 |
| 1/29/19 | 2018-2019 | Londo | -22.99 | 1.47 | 5 |
| 1/29/19 | 2018-2019 | Martinson | -23.36 | 1.15 | 30 |
| 2/4/19 | 2018-2019 | Londo | -23.16 | 1.85 | 5 |
| 2/4/19 | 2018-2019 | Martinson | -24.16 | 2.34 | 30 |
| 2/11/19 | 2018-2019 | Londo | -21.34 | 0.71 | 10 |
| 2/12/19 | 2018-2019 | Martinson | -23.05 | 2.17 | 30 |
| 2/18/19 | 2018-2019 | Martinson | -24.01 | 1.91 | 30 |
| 2/21/19 | 2018-2019 | Londo | -22.58 | 0.73 | 5 |
| 2/26/19 | 2018-2019 | Martinson | -24.51 | 3.03 | 30 |
| 2/28/19 | 2018-2019 | Londo | -22.02 | 1.99 | 5 |
| 3/4/19 | 2018-2019 | Martinson | -24.25 | 2.54 | 30 |
| 3/7/19 | 2018-2019 | Londo | -20.90 | 2.12 | 5 |
| 3/12/19 | 2018-2019 | Martinson | -26.32 | 3.91 | 30 |
| 3/13/19 | 2018-2019 | Londo | -21.81 | 1.48 | 10 |
| 3/18/19 | 2018-2019 | Martinson | -23.94 | 1.80 | 30 |
| 3/20/19 | 2018-2019 | Londo | -19.57 | 0.48 | 5 |
| 3/26/19 | 2018-2019 | Martinson | -25.42 | 3.18 | 30 |
| 3/28/19 | 2018-2019 | Londo | -19.87 | 1.00 | 3 |
| 4/2/19 | 2018-2019 | Martinson | -21.89 | 1.88 | 30 |
| 4/3/19 | 2018-2019 | Londo | -19.02 | 0.93 | 3 |
| 4/10/19 | 2018-2019 | Londo | -14.21 | 0.84 | 3 |
| 4/16/19 | 2018-2019 | Londo | -8.68 | 0.55 | 4 |

| | | | | | | |
|---|---|---|---|---|---|---|
| 9/12/19 | 2019-2020 | Londo | -6.71 | | 0.67 | 4 |
| 9/24/19 | 2019-2020 | Londo | -8.42 | | 1.22 | 8 |
| 10/2/19 | 2019-2020 | Londo | -9.61 | | 0.75 | 5 |
| 10/10/19 | 2019-2020 | Londo | -9.27 | | 1.20 | 5 |
| 10/18/19 | 2019-2020 | Londo | -11.66 | | 2.17 | 10 |
| 10/23/19 | 2019-2020 | Londo | -11.36 | | 1.47 | 9 |
| 10/30/19 | 2019-2020 | Londo | -12.61 | | 0.90 | 8 |
| 11/6/19 | 2019-2020 | Londo | -13.30 | | 1.47 | 10 |
| 11/13/19 | 2019-2020 | Londo | -17.66 | | 1.02 | 9 |
| 11/20/19 | 2019-2020 | Londo | -19.36 | | 1.10 | 10 |
| 11/27/19 | 2019-2020 | Londo | -19.65 | | 0.45 | 16 |
| 12/4/19 | 2019-2020 | Londo | -21.00 | | 1.70 | 10 |
| 12/11/19 | 2019-2020 | Londo | -20.02 | | 2.45 | 6 |
| 12/18/19 | 2019-2020 | Londo | -18.89 | | 2.14 | 6 |
| 12/19/19 | 2019-2020 | Londo | -22.46 | | 0.33 | 5 |
| 12/24/19 | 2019-2020 | Londo | -21.51 | | 1.22 | 7 |
| 1/2/20 | 2019-2020 | Londo | -21.99 | | 0.97 | 10 |
| 1/7/20 | 2019-2020 | Londo | -22.38 | | 0.82 | 7 |
| 1/15/20 | 2019-2020 | Londo | -24.35 | | 2.55 | 7 |
| 1/22/20 | 2019-2020 | Londo | -22.36 | | 1.29 | 7 |
| 1/29/20 | 2019-2020 | Londo | -22.06 | | 0.73 | 8 |
| 2/5/20 | 2019-2020 | Londo | -21.46 | | 1.65 | 7 |
| 2/11/20 | 2019-2020 | Londo | -22.58 | | 1.42 | 7 |
| 2/12/20 | 2019-2020 | Londo | -23.13 | | 1.82 | 7 |
| 2/19/20 | 2019-2020 | Londo | -24.10 | | 0.34 | 3 |
| 2/26/20 | 2019-2020 | Londo | -20.68 | | 2.76 | 6 |
| 3/4/20 | 2019-2020 | Londo | -21.24 | | 1.75 | 13 |
| 3/11/20 | 2019-2020 | Londo | -19.50 | | 1.53 | 2 |
| 3/18/20 | 2019-2020 | Londo | -17.09 | | 0.68 | 3 |
| 3/25/20 | 2019-2020 | Londo | -16.73 | NA | | 1 |
| 10/2/20 | 2020-2021 | Londo | -9.57 | | 1.21 | 8 |
| 10/8/20 | 2020-2021 | Londo | -10.26 | | 1.32 | 10 |
| 10/14/20 | 2020-2021 | Londo | -11.49 | | 0.62 | 8 |
| 10/22/20 | 2020-2021 | Londo | -10.49 | | 1.07 | 5 |
| 10/29/20 | 2020-2021 | Londo | -11.05 | | 1.85 | 8 |
| 11/4/20 | 2020-2021 | Londo | -13.44 | | 3.26 | 7 |
| 11/10/20 | 2020-2021 | Londo | -14.24 | | 0.71 | 6 |
| 11/18/20 | 2020-2021 | Londo | -18.61 | | 1.14 | 16 |
| 11/25/20 | 2020-2021 | Londo | -18.41 | | 1.04 | 12 |
| 12/2/20 | 2020-2021 | Londo | -19.05 | | 1.82 | 7 |

| | | | | | |
|---|---|---|---|---|---|
| 12/10/20 | 2020-2021 | Londo | -21.35 | 1.61 | 10 |
| 12/15/20 | 2020-2021 | Londo | -19.57 | 1.80 | 7 |
| 12/23/20 | 2020-2021 | Londo | -21.01 | 1.06 | 10 |
| 12/30/20 | 2020-2021 | Londo | -23.59 | 1.04 | 8 |
| 1/6/21 | 2020-2021 | Londo | -21.00 | 1.27 | 8 |
| 1/13/21 | 2020-2021 | Londo | -23.46 | 1.36 | 9 |
| 1/20/21 | 2020-2021 | Londo | -23.60 | 1.79 | 7 |
| 1/27/21 | 2020-2021 | Londo | -25.42 | 0.53 | 9 |
| 2/3/21 | 2020-2021 | Londo | -20.40 | 2.15 | 6 |
| 2/10/21 | 2020-2021 | Londo | -20.65 | 4.21 | 11 |
| 2/17/21 | 2020-2021 | Londo | -23.11 | 1.55 | 10 |
| 2/24/21 | 2020-2021 | Londo | -23.50 | 3.02 | 9 |
| 3/3/21 | 2020-2021 | Londo | -22.82 | 1.08 | 9 |
| 3/10/21 | 2020-2021 | Londo | -22.19 | 1.79 | 8 |
| 3/17/21 | 2020-2021 | Londo | -19.65 | 1.29 | 11 |
| 3/24/21 | 2020-2021 | Londo | -18.40 | 0.96 | 7 |
